# AvrE1 and HopR1 from *Pseudomonas syringae* pv. *actinidiae* are additively required for full virulence on kiwifruit

**DOI:** 10.1101/2020.06.18.158634

**Authors:** Jay Jayaraman, Minsoo Yoon, Emma R. Applegate, Erin A. Stroud, Matthew D. Templeton

## Abstract

*Pseudomonas syringae* pv. *actinidiae* ICMP 18884 biovar 3 (*Psa3*) produces necrotic lesions during infection of its kiwifruit host. Bacterial growth *in planta* and lesion formation are dependent upon a functional type III secretion system (T3S), which translocates multiple effector proteins into host cells. Associated with the T3S locus is the conserved effector locus (CEL), which has been characterised and shown to be essential for the full virulence in other *P. syringae* pathovars. Two effectors at the CEL, *hopM1* and *avrE1*, as well as an *avrE1*-related non-CEL effector, *hopR1*, have been shown to be redundant in the model pathogen *P. syringae* pv. *tomato* DC3000 (*Pto*), a close relative of *Psa*. However, it is not known whether CEL-related effectors are required for *Psa* pathogenicity. The *Psa3* allele of *hopM1*, and its associated chaperone, *shcM*, have diverged significantly from their orthologs in *Pto*. Furthermore, the CEL effector *hopAA1-1*, as well as a related non-CEL effector, *hopAA1-2*, have both been pseudogenised. We have shown that HopM1 does not contribute to *Psa3* virulence due to a truncation in *shcM*, a truncation conserved in the *Psa* lineage, likely due to the need to evade HopM1-triggered immunity in kiwifruit. We characterised the virulence contribution of CEL and related effectors in *Psa3* and found that only *avrE1* and *hopR1*, additively, are required for *in planta* growth and lesion production. This is unlike the redundancy described for these effectors in *Pto* and indicates that these two *Psa3* genes are key determinants essential for kiwifruit bacterial canker disease.

## Introduction

*Pseudomonas syringae* is a model plant pathogen for studying bacterial pathogenicity, in particular, the molecular mechanisms of disease establishment. The *P. syringae* species complex is currently represented by 64 pathovars, with each pathovar identified as infecting a select host plant genus, but often possessing the capacity to infect more, being restricted by plant immunity and environmental parameters (Berge et al., 2014; Laflamme et al., 2020; Morris et al., 2019; Xin et al., 2016). Collectively, the pathovars of *P. syringae* cause disease on the vast majority of agriculturally important crop plants, demonstrating the ubiquity and flexibility of this species globally. Regular disease outbreaks, caused by emergent *P. syringae* strains, frequently threaten global agricultural productivity (Xin et al., 2018).

Comparative and population genomics studies of isolates collected from agricultural and environmental sources have transformed our understanding of diseases caused by *P. syringae* (Dillon et al., 2019b, 2019a; McCann et al., 2017). The *P. syringae* species complex is divided into 13 phylogroups on the basis of multilocus sequence analysis, with the three major pathogenic phylogroups (numbered 1–3) with their core genomes forming distinct lineages, but with accessory genetic components shared between them through horizontal gene transfer (Berge et al., 2014; Dillon et al., 2019b). A near-universally common genetic component of these plant pathogens is a canonical tripartite pathogenicity island composed of a *hrp* and *hrc* gene cluster encoding a type III secretion system (T3S), flanked by the conserved effector locus (CEL), and the exchangeable effector locus (EEL) (Xin et al., 2018). The T3S delivers bacterial effector proteins (T3Es) into host cells and is necessary for pathogenesis (Büttner and He, 2009; Clarke et al., 2010). While effectors encoded by the EEL vary between pathovars and strains, with several expected to be functionally redundant, the CEL encodes between two and four highly conserved effector genes: *hopAA1-1, hopM1, avrE1*, and occasionally *hopN1* (Deng et al., 2003).

Evolutionarily conserved CEL-encoded T3Es are required for (and predictive of) pathogenicity (Alfano et al., 2000; Baltrus et al., 2011; Dillon et al., 2019b). There is evidence of redundancy in this T3E group, but studies of this phenomenon as well as associated contributions of each CEL effector are limited to the type strain *P. syringae* pv. *tomato* (*Pto*) DC3000. These studies have revealed functional redundancy between CEL T3Es *hopM1* and *avrE1*, and an *avrE*-related non-CEL T3E, *hopR1*, in a host species-dependent manner (Badel et al., 2006; Kvitko et al., 2009). Whether these results are representative of other *P. syringae* strains on their respective hosts is currently unknown.

In this study, the CEL locus T3Es and their non-CEL homologs/orthologs were characterised from the kiwifruit bacterial canker pathogen, *P. syringae* pv. *actinidiae* (*Psa*), and compared and contrasted with orthologs from *Pto* DC3000. *Psa* and *Pto* strains are closely related in *P. syringae* phylogroup 1 (PG1), albeit from different clades, PG1a and PG1b, respectively (Berge et al., 2014). *Psa* HopM1 was found to be non-functional due to truncation of its chaperone ShcM, while AvrE1 and HopR1 were found to be functional and non-redundantly required for full virulence in *Psa* infection of kiwifruit.

## Results

### The CEL is required for pathogenicity of *Psa3* in *Actinidia chinensis* var. *chinensis* ‘Hort16A’

The CEL of *Psa* biovar 3 ICMP 18884 (hereafter *Psa3* V13) was identified previously during genome annotation via the NCBI Prokaryotic Genome Annotation Pipeline (PGAP) (Templeton et al., 2015), and comprises T3E genes *hopN1* (with its chaperone *shcN*), *hopAA1-1*, the T3S helper gene *hrpW1, hopM1* (with its chaperone *shcM*), and *avrE1* (with its chaperone *shcE*) (Fig. 1A). It is highly similar to and syntenic with the CEL of type strain *Pto* DC3000 (Table 1). In *Psa3*, however, there is a 14-bp indel in the genomic sequence of *hopAA1-1*, resulting in an early truncation of its peptide sequence and thus poor similarity in this effector (Badel et al., 2006). To investigate whether the CEL in *Psa3* V13 has an essential role in pathogenicity a knockout of the entire 14.2 kb region of the CEL was generated (*ΔCEL*). We assessed ‘pathogen fitness’ by measuring *in planta* bacterial growth and ‘virulence’ by examining disease symptom development. Pathogenicity testing of the *ΔCEL* mutant was carried out on *Actinidia chinensis* var. *chinensis* ‘Hort16A’ plantlets along with wildtype *Psa3* V13 and the T3S-deficient *ΔhrcC* strain. Loss of the CEL resulted in significant reduction in *Psa3* V13 fitness, as assessed by *in planta* growth at both 6-and 12-days post-inoculation (dpi) (Fig. 1B), but not to the extent of that observed for the type III secretion system *ΔhrcC* mutant. Reflecting the reduction of *in planta* growth was a significant reduction in both leaf lesion development and plant death in ‘Hort16A’ plants at 50 dpi, compared to the wildtype strain, indicating an associated reduction in virulence (Fig. 1C). Taken together, these results suggest that the CEL is required for full virulence of *Psa3* on ‘Hort16A’ kiwifruit plants.

**Table 1.**
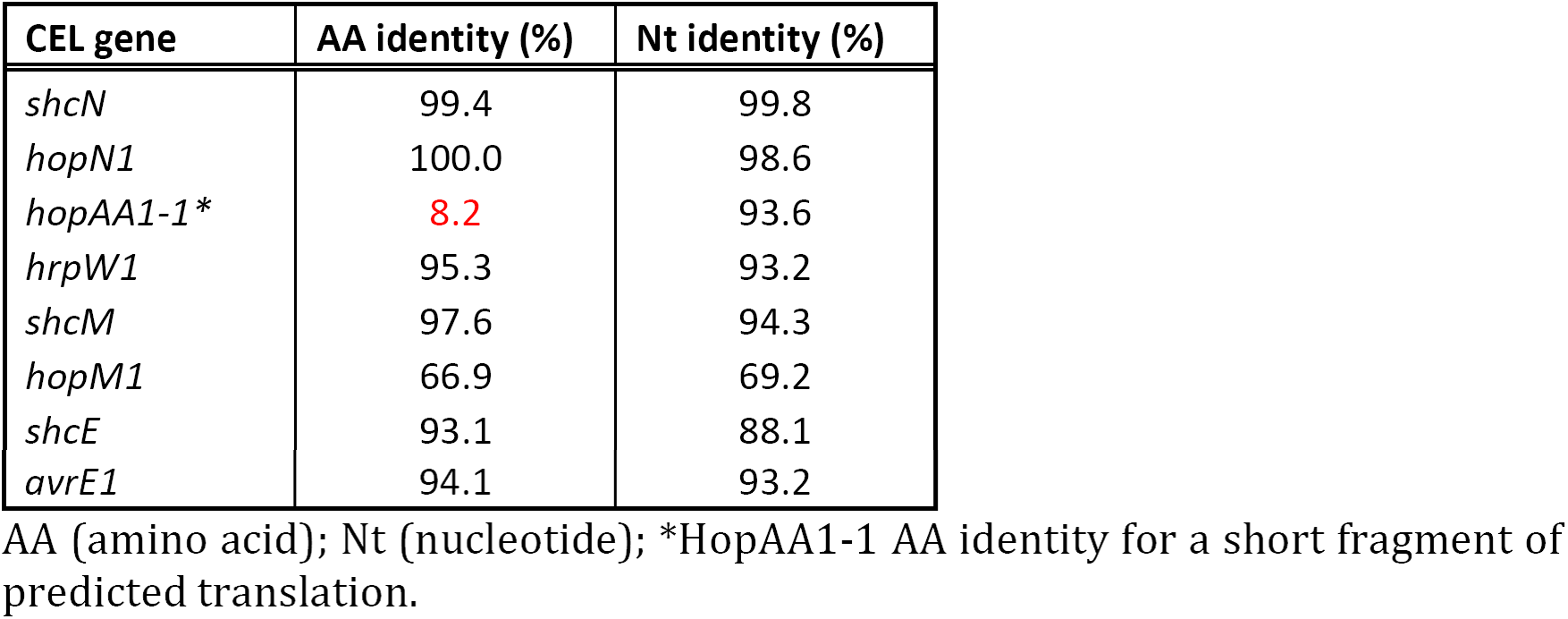
Comparison of conserved effector locus (CEL) encoded genes from *Pseudomonas syringae* pv. *actinidiae* ICMP 18884 biovar 3 (*Psa3*) V13 and *P. syringae* pv. *tomato* (*Pto*) DC3000.

| CEL gene | AA identity (%) | Nt identity (%) |
| --- | --- | --- |
| *shcN* | 99.4 | 99.8 |
| *hopN1* | 100.0 | 98.6 |
| *hopAA1-1\** | 8.2 | 93.6 |
| *hrpW1* | 95.3 | 93.2 |
| *shcM* | 97.6 | 94.3 |
| *hopM1* | 66.9 | 69.2 |
| *shcE* | 93.1 | 88.1 |
| *avrE1* | 94.1 | 93.2 |

AA (amino acid); Nt (nucleotide); \*HopAA1-1 AA identity for a short fragment of predicted translation.

**Figure 1.**
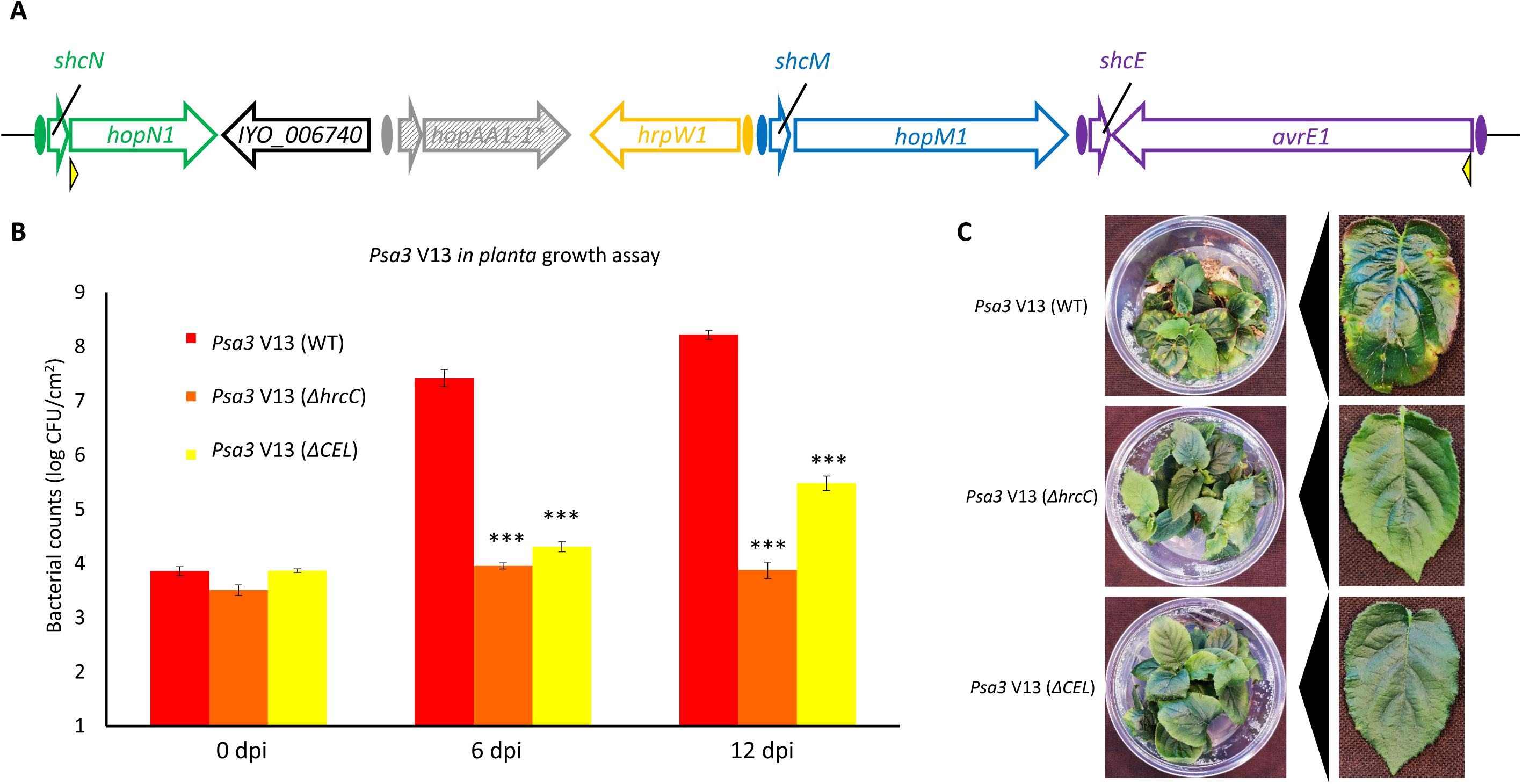
The conserved effector locus in *Pseudomonas syringae* pv. *actinidiae* ICMP 18884 biovar 3 (*Psa3*) is required for full virulence in *Actinidia chinensis* var. *chinensis* ‘Hort16A’ kiwifruit plants. **(A)** Overview schematic of the conserved effector locus (CEL) region and open reading frames (ORFs) in *Psa3*. ORFs (arrows) indicate the direction of transcription, with ovals denoting the presence of an HrpL box. The asterisk and hatched shading for the *hopAA1-1* gene indicates the truncation of the original gene to generate a two-part pseudogene. The two yellow arrows indicate the region deleted in the *Psa3 ΔCEL* mutant. **(B)** *P*sa ICMP 18884 (*Psa3* V13) growth in ‘Hort16A’ plants. *Psa3* V13 (wildtype, WT), *ΔhrcC* or *ΔCEL* mutants were flood-inoculated at ∼10^6^ colony forming units; CFU/mL into ‘Hort16A’ leaves, and bacterial growth was determined 0, 6, and 12 days post-inoculation (dpi). Error bars represent standard error of the mean from four pseudobiological replicates. Asterisks indicate results of a two-tailed Student’s *t*-test between the selected sample and wildtype *Psa3* V13; ***(*P* < 0.001). The experiment was conducted four times with similar results. **(C)** Symptom development in ‘Hort16A’ leaves at 50 dpi for strains inoculated in (B). Photographs taken of a single representative pottle and a representative leaf displaying typical disease lesion symptoms, if present.

### AvrE1 is responsible for CEL-conferred virulence of *Psa3*

Having established a significant role for the CEL in the pathogenicity of *Psa3* V13, the contribution of individual CEL-encoded effectors to *Psa3* pathogenicity was investigated. Broad host-range plasmid-borne copies of each functional effector gene from the CEL were transformed into the *Psa3* V13 *ΔCEL* mutant (Fig. 2A). *In planta* bacterial growth assays in ‘Hort16A’ plantlets revealed that only plasmid-borne *avrE1* (p.*avrE1*) significantly restored *in planta* growth to the *ΔCEL* mutant (Fig. 2B), but *Psa3* V13 *ΔCEL* + p.*avrE1* was reduced compared to wildtype at 12 dpi. This difference was not due to plasmid loss during *in planta* growth (Fig. S1). None of the plasmid-borne CEL effectors were able to fully restore virulence as demonstrated by the lack of lesion development and plant death at 50 dpi, however (Fig. 2C).

**Figure 2.**
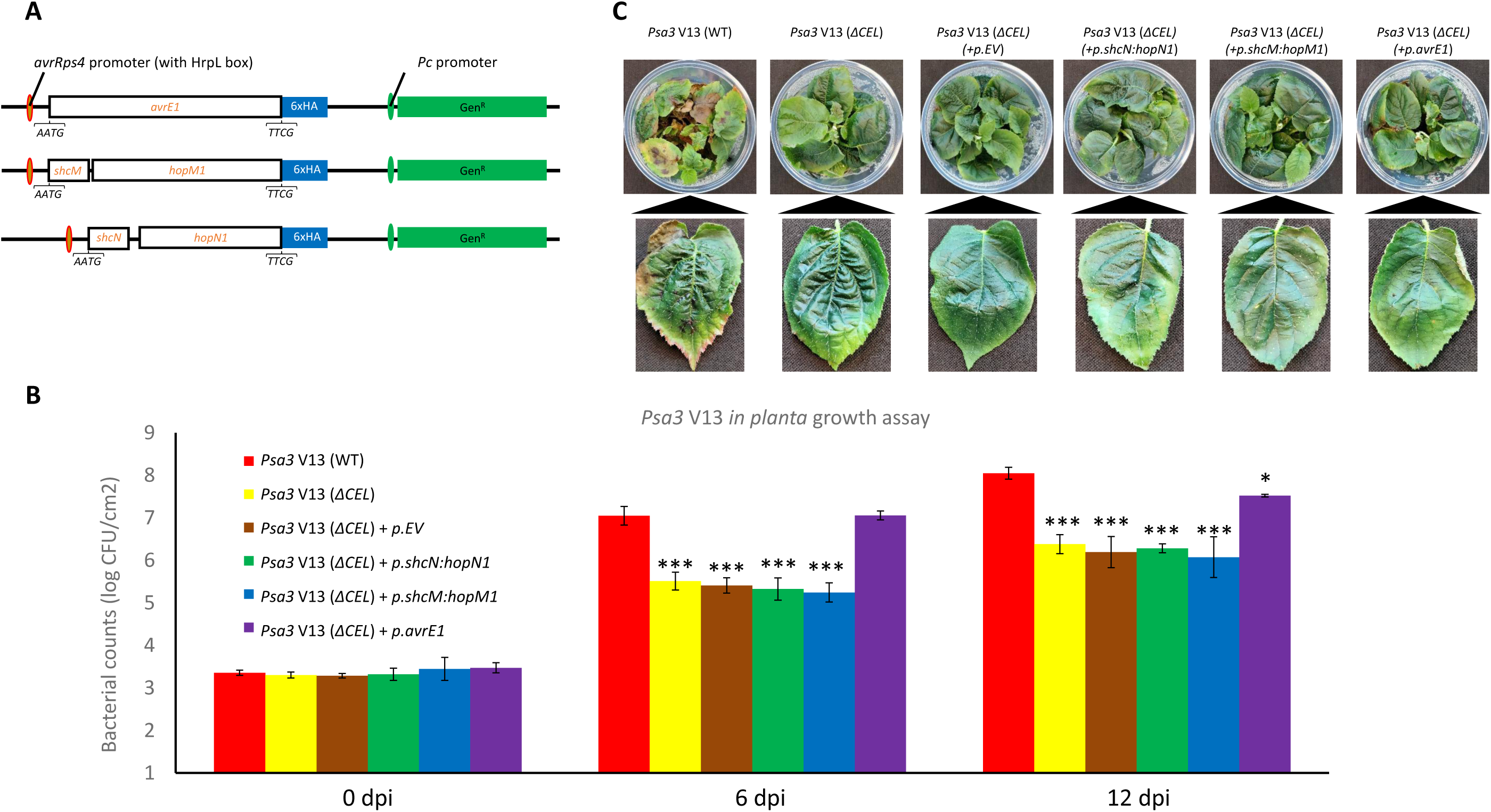
Plasmid-borne *avrE1* is able to largely restore virulence to *Pseudomonas syringae* pv. *actinidiae* ICMP 18884 biovar 3 (*Psa3*) *ΔCEL*. **(A)** Schematic of pBBR1MCS-5 vector constructs (Jayaraman et al., 2017) for under the AvrRps4 HrpL box for *hopN1* with chaperone *shcN, hopM1* with chaperone *shcM*, or *avrE1* without its chaperone. The effectors were tagged with 6xHA, and plasmids were selected for by a gentamicin resistance cassette under a strong *Pc* promoter. The 5’ (AATG; including the first codon of the chaperone/effector) and 3’ (TTCG; non-native sequence) overhangs used for assembly of the final construct are indicated. **(B)** *Psa3* V13 (wildtype, WT), conserved effector locus mutant (*ΔCEL*), *ΔCEL* carrying empty vector (p.EV), or *ΔCEL* plasmid complementation mutants, were flood-inoculated at ∼10^6^ colony forming units; CFU/mL into *Actinidia chinensis* var. *chinensis* ‘Hort16A’, leaves and bacterial growth was determined 0, 6, and 12 days post-inoculation (dpi). Error bars represent standard error of the mean from four pseudobiological replicates. Asterisks indicate results of a two-tailed Student’s *t*-test between the selected sample and wildtype (*Psa3* V13); *(*P* < 0.05), ***(*P* < 0.001). The experiment was conducted three times with similar results. **(C)** Symptom development in ‘Hort16A’ leaves at 50 dpi for strains inoculated in (B). Photographs taken of a single representative pottle and a representative leaf displaying typical disease lesion symptoms, if present.

The lack of full complementation by p.*avrE1* may in part be explained by either the absence of its chaperone *shcE* (Lorang and Keen, 1995) or an unknown effect of genomic localisation for *avrE1* (and/or *hopM1)* required for appropriate levels of expression and/or secretion. Alternatively, T3S helper *hrpW1*, which was not included in the original screen, may be involved in conferring full virulence. This gene is expressed at high levels during *in planta* infection of *Psa3* compared to other functionally redundant helpers (Fig. S2) (Kvitko et al., 2007; McAtee et al., 2018). To assess this, genomic knock-in constructs of *hrpW1, hopM1* (with chaperone *shcM*), or *avrE1* (with chaperone *shcE*) were generated with their native HrpL box promoters and reintegrated into *Psa3* V13 genome at the CEL knock-out site (Fig. 3A). Assessment of *in planta* bacterial growth confirmed our previous result that *avrE1* restored *in planta* growth to the *ΔCEL* mutant, and further, that inclusion of *shcE*, restored higher growth at both 6 and 12 dpi (Fig. 3B). In addition, the genomic *shcE:avrE1* knock-in fully restored symptomology with leaf lesions and plant death observed at 50 dpi, suggesting virulence at this locus is largely conferred by AvrE1 (Fig. 3C). In contrast, genomic knock-in of *hopM1* with chaperone *shcM* (*shcM:hopM1*) did not restore *in planta* growth or virulence to the *Psa3 ΔCEL* mutant.

**Figure 3.**
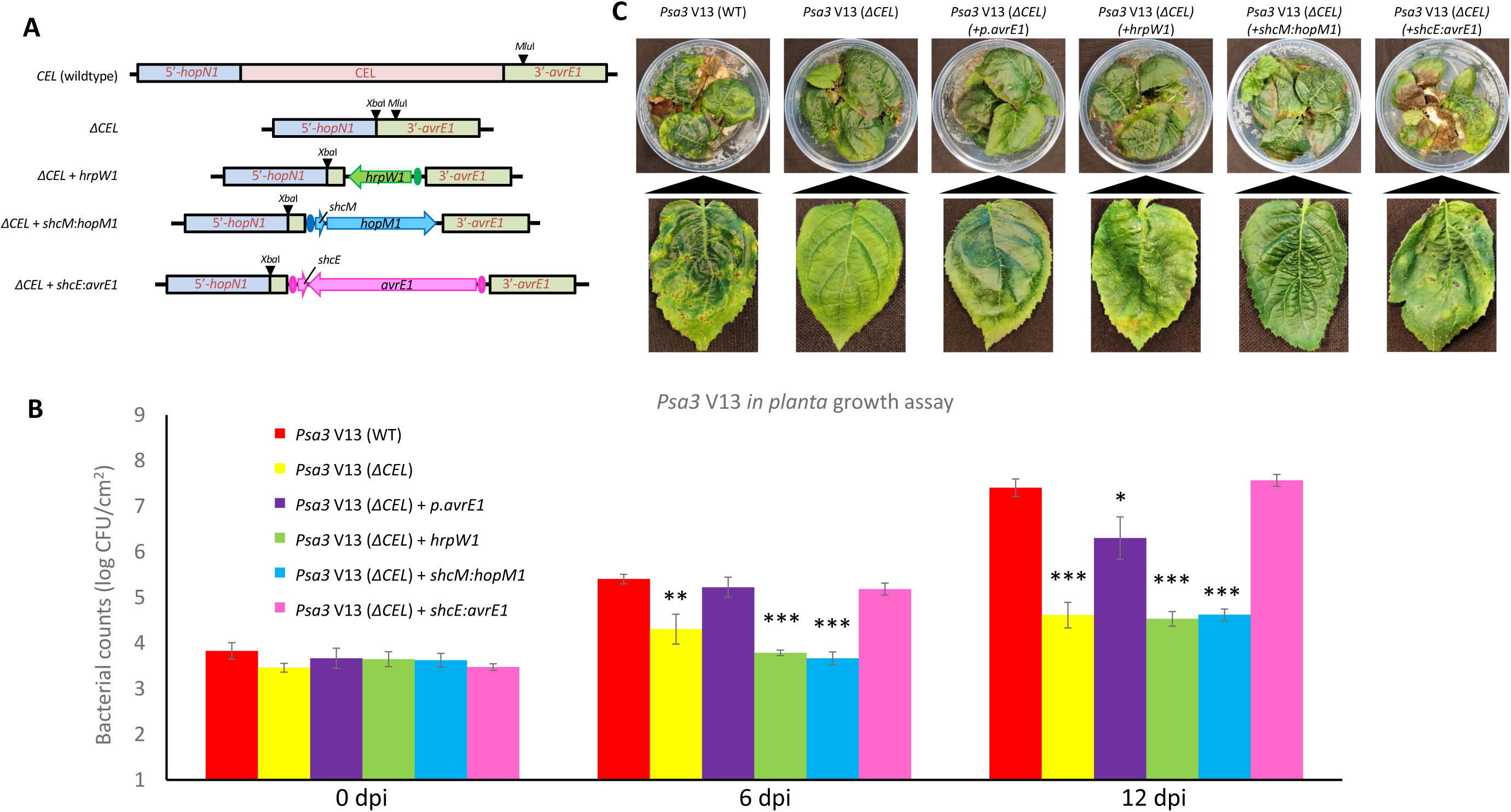
*avrE1* genomic knock-in with *shcE* fully restores virulence to *Pseudomonas syringae* pv. *actinidiae* ICMP 18884 biovar 3 (*Psa3*) *ΔCEL*. **(A)** Schematic of the conserved effector locus (CEL) locus (wildtype), knockout with introduced *Xba*I site (*ΔCEL*), knock-in for *hrpW1* (*ΔCEL+hrpW1*), knock-in for *hopM1* with chaperone *shcM* (*ΔCEL+shcM:hopM1*), or knock-in for *avrE1* with chaperone *shcE* (*ΔCEL+shcE:avrE1*). The knock-in was made at the native *Mlu*I site indicated in the 3’ region of the *ΔCEL* knockout construct. Coloured ovals indicate inclusion of the relevant HrpL binding site promoters included in the knock-in constructs. **(B)** *Psa3* V13 (wildtype, WT), *ΔCEL* mutant, *ΔCEL* with *avrE1* plasmid-complemented mutant, or *ΔCEL* with genomic knock-in *hrpW1, shcM:hopM1*, or *shcE:avrE1* were inoculated by flooding at ∼10^6^ CFU/mL into *Actinidia chinensis* var. *chinensis* ‘Hort16A’ leaves, and bacterial growth was determined 0, 6, and 12 days post-inoculation (dpi). Error bars represent standard error of the mean from four pseudobiological replicates. Asterisks indicate results of a two-tailed Student’s *t*-test between the selected sample and wildtype (*Psa3* V13); *(*P* < 0.05), **(*P* < 0.01), ***(*P* < 0.001). The experiment was conducted three times with similar results. **(C)** Symptom development in ‘Hort16A’ leaves at 50 dpi for strains inoculated in (B). Photographs taken of a single representative pottle and a representative leaf displaying typical disease lesion symptoms, if present.

### HopM1 function in *Psa3* is affected by truncation of its chaperone ShcM

The *shcM:hopM1* genomic knock-in was unable to complement the *Psa3 ΔCEL* mutant, despite *Psa3* having a full length *hopM1* allele (Fig. 4) (Templeton et al., 2015). Closer examination of the *Psa hopM1* genomic region (and its related alleles in *P. syringae* phylogroup 1), including *shcM*, suggested that the *Psa* lineage lacks the ability to deliver a fully functional HopM1 due to an early truncation in either *shcM* (*P. syringae* pv. *morsprunorum* (*Pmp*) and *Psa1-6*) or *hopM1* (*P. syringae* pv. a*ctinidifoliorum*: *Pfm*; previously *Psa4*) itself (Fig. 4). All *Psa* biovars shared a single nucleotide insertion in the associated chaperone *shcM* that is unique to *Psa. Pmp* possesses a single nucleotide deletion in its *shcM* sequence, and *Pfm* is truncated by a small nucleotide deletion in *hopM1. Psa3* strains possess an additional single nucleotide deletion downstream of this aforementioned deletion, while *Psa6* has an insertion downstream, but these are silent mutations since the upstream deletion inserts a stop codon into the coding sequence.

**Figure 4.**
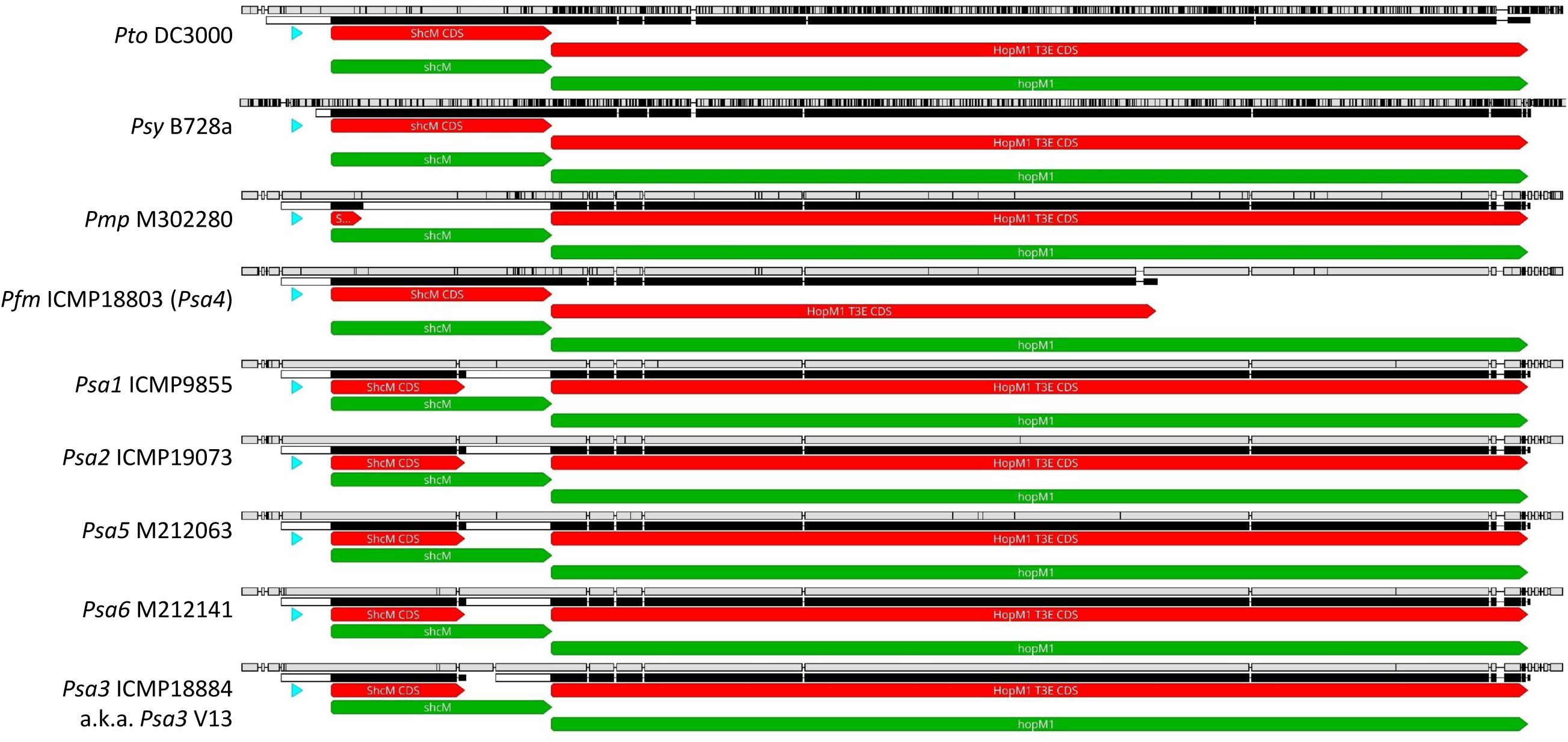
Strains from the *Pseudomonas syringae* pv. *actinidiae* ICMP 18884 biovar 3 (*Psa3*) lineage have consistently disrupted *shcM* or *hopM1*. Nucleotide sequence alignment of the *shcM*:*hopM1* regions of *Pto* DC3000 (set as reference), *Psy* B728A, *Pmp* M302280, and representative members of *Psa* biovars (*Psa1-6*), respectively: *Psa1* ICMP 9855, *Psa2* ICMP 19073, *Psa3* ICMP 18884, *Pfm* ICMP 18803 (previously *Psa4*), *Psa5* MAFF 212063, and *Psa6* MAFF 212141, as viewed in Geneious software. The top bar (black/white) in each panel represents nucleotide sequence similarity, with the bar underneath representing amino acid similarity. Red arrows indicate coding sequences, and green arrows indicate genomic regions (ORFs). The HrpL box for the *shcM*:*hopM1* operon is represented by the blue arrowhead for each sequence.

To assess whether HopM1 delivery was affected by the truncation in the ShcM protein, plasmid-borne versions of each of the CEL effectors (with their chaperones, if available) were transformed into wildtype *Pseudomonas fluorescens* (*Pfo*) Pf0-1, or an artificially generated T3S-carrying *P. fluorescens* strain, *Pfo* Pf0-1(T3S) (Thomas et al., 2009), and infiltrated into *Nicotiana benthamiana* leaves for protein expression (Fig. 5) or cell death analysis (Fig. S4). To avoid the effects of differences in promoter sequences affecting expression, *shcM* from *Pto* DC3000 (*shcM*_*Pto*_) was cloned under the same *avrRps4* promoter to control expression of the plasmid-borne *hopM1* operon used to assess *shcM*_*Psa*_ described earlier (Jayaraman et al., 2017). *shcM*_*Pto*_ was cloned as a *cis*-complementation construct ahead of *hopM1*_*Psa*_ and transformed into *Pfo* Pf0-1 or *Pfo* Pf0-1(T3S) strains for comparison.

**Figure 5.**
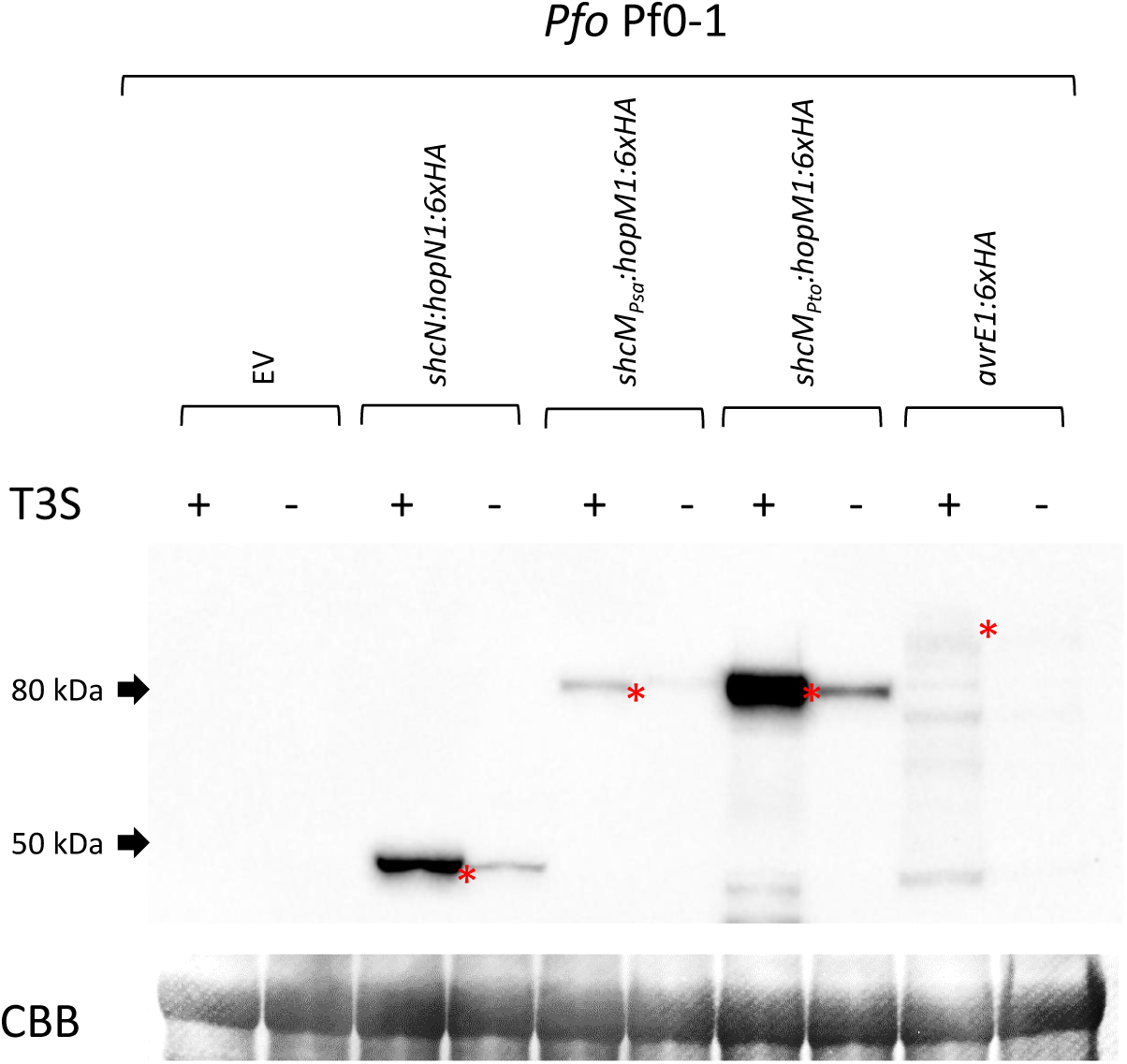
*Pseudomonas fluorescens* (*Pfo*) Pf0-1 carrying *shcM* from *P. syringae* pv. *tomato* (*Pto*) DC3000 shows increased *in planta* secretion of HopM1 in a type III secretion system (T3S)-dependent manner. *In planta* secretion assay for empty vector (EV), HopN1 (vector with shcN), HopM1 (vector with *shcM*_*Psa*_ or *shcM*_*Pto*_), or AvrE1 (no chaperone) proteins tagged with 6xHA from *Pfo* Pf0-1 (T3S +) or *Pfo* Pf0-1 (wildtype, T3S -). *Pfo* strains were infiltrated at Abs_600_ = 1.5 into *Nicotiana benthamiana* plants and leaf samples harvested at 6 h post-infiltration, protein extracted and western blots conducted using α–HA antibody. Red asterisks indicate expected sizes for each tagged protein band. The Coomassie Brilliant Blue (CBB)-stained band for Rubisco is presented as loading control.

Interestingly, at higher bacterial loads (OD_600_ = 1), AvrE1 (without chaperone ShcE) and HopM1 (with native chaperone ShcM_*Psa*_) were both able to trigger a cell death response in *N. benthamiana*. However, the cell death triggered by HopM1 was weak/sporadic (Fig. S3A), suggesting poor expression or delivery. This was confirmed by *in planta* effector secretion assays (Fig. 5). In contrast, *cis*-complementation with full length *shcM*_*Pto*_ along with *hopM1* in Pf0-1(T3S) was able to confer robust HopM1-triggered cell death even at reduced bacterial loads (Fig. S3B-C) and improved secretion of HopM1 (Fig. 5).

### HopM1 triggers immunity in *A. chinensis* ‘Hort16A’

To determine whether *shcM* complementation restored HopM1-mediated virulence, the *Psa3* V13 (wildtype), *Psa3 ΔCEL* mutant, *Psa3 ΔCEL* + *hopM1* (genomic knock-in of *shcM*_*Psa*_:*hopM1*), and *Psa3 ΔCEL* + *avrE1* (genomic knock-in of *shcE:avrE1*) strains were transformed with plasmid-borne *shcM*_*Pto*_ (*trans*-complemented), and assessed by *in planta* bacterial growth assays in ‘Hort16A’ plantlets. Intriguingly, while *shcM*_*Pto*_ complementation did not restore virulence, it consistently reduced the *in planta* growth of *Psa3* V13 (wildtype) and *Psa3 ΔCEL* + *hopM1*, but not of *Psa3 ΔCEL* and *Psa3 ΔCEL* + *avrE1* suggesting that HopM1 triggers an immune response (Fig. S4A). This difference was not due to plasmid loss during *in planta* growth (Fig. S4B).

To further confirm the contribution of the *shcM* allele in triggering immunity in ‘Hort16A’ plants, *cis*-complementation by the *shcM*_*Pto*_ construct alongside the original *shcM*_*Psa*_ construct in *Psa3* V13 wildtype, *Psa3 ΔCEL*, and *Psa3 ΔCEL* + *avrE1* (genomic knock-in of *shcE*:*avrE1*) was tested. When transformed with plasmid-borne *shcM*_*Pto*_, a consistent reduction of *in planta* growth confirmed that the restoration of HopM1 delivery by *Psa3* reduces *in planta* growth in a HopM1-dependent manner (Fig. 6A). Despite this finding, no changes in *shcM*-dependent virulence were detected as determined by a lack of reduction in disease symptoms (Fig. 6B). This decrease in *in planta* growth was accompanied by an associated an increase in *AcWRKY70a* defense gene expression in ‘Hort16A’ plants at 24 h post-inoculation (Fig. 6C). Notably, the *cis*-complementation of *shcM*_*Pto*_ allowed for increased *in vitro* secretion of HopM1 in a T3S-dependent manner, suggesting that the ShcM_*Pto*_-mediated increase in secretion of HopM1 facilitated this change in response (Fig. S5). Therefore, the HopM1-dependent reduction of *in planta* growth and associated defense gene induction indicates that HopM1 may trigger immunity in ‘Hort16A’ plants, and suggests that the loss of ShcM function through truncation is a mechanism of reducing HopM1-triggered immunity in its host kiwifruit plants.

**Figure 6.**
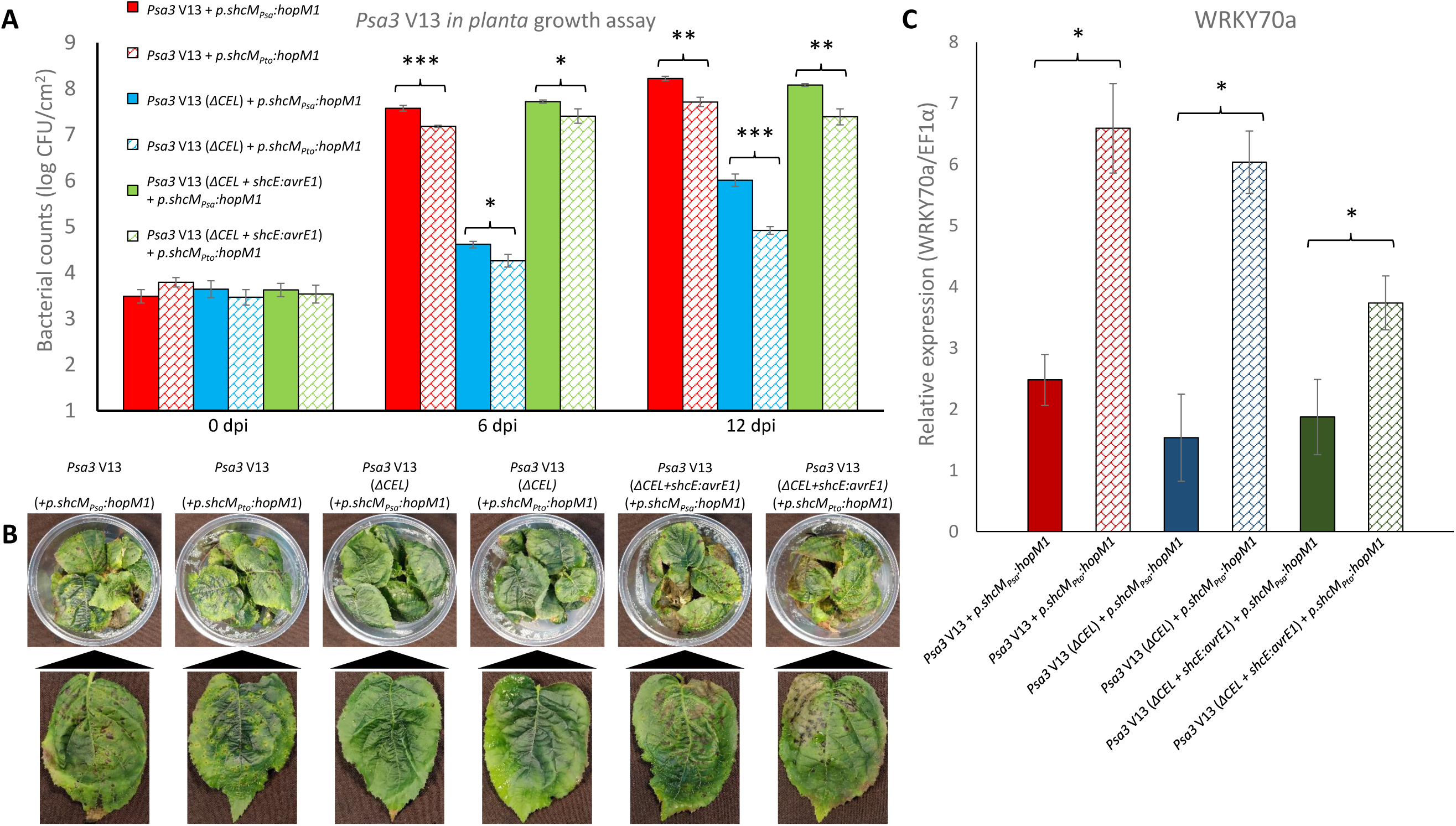
*Pseudomonas syringae* pv. *actinidiae* ICMP 18884 biovar 3 (*Psa3*) carrying *shcM* from *P. syringae* pv. *tomato* (*Pto*) DC3000 is reduced in host colonization but not virulence in a *hopM1*-dependent manner. **(A)** *Psa3* V13 (wildtype), *ΔCEL* mutant, or *ΔCEL* with genomic knock-in *shcE:avrE1*, with or without plasmid-borne *shcM*_*Psa*_:*hopM1* or *shcM*_*Pto*_:*hopM1*, were inoculated by flooding at ∼10^6^ colony forming units; CFU/mL on *Actinidia chinensis* var. *chinensis* ‘Hort16A’, and bacterial growth was determined 0, 6, and 12 days post-inoculation (dpi). Error bars represent standard error of the mean from four pseudobiological replicates. Asterisks indicate results of a two-tailed Student’s *t*-test between selected samples indicated; *(*P* < 0.05), **(*P* < 0.01), ***(*P* < 0.001). The experiment was conducted three times with similar results. **(B)** Symptom development in ‘Hort16A’ at 50 dpi for strains inoculated in (A). Photographs taken of a single representative pottle and a representative leaf displaying typical disease lesion symptoms, if present. **(C)** Defense gene expression in on ‘Hort16A’ in response to shcM complementation. *Psa3* strains from (A) were -inoculated by flooding at ∼10^7^ CFU/mL and samples taken at 24 h post-inoculation and defense gene (*AcWRKY70a*; Acc03057.1) expression determined from extracted RNA by quantitative polymerase chain reaction. Expression for *AcWRKY70a* is normalised to internal *EF1α* expression. Error bars indicate standard error from three pseudobiological replicates. Asterisks indicate results of Student’s t-test between selected samples; *(*P* < 0.05).

Comparison of the *hopM1* alleles for type strains from phylogroups 1, 2 and 3 suggests that the *Psa* lineage (*Psa, Pfm, Pmp, P. syringae* pv. *theae* (*Pth*), and *P. syringae* pv. *avellenae* (*Pav*)), a subset of clade 1b in phylogroup 1, possesses a *hopM1* allele that has a distinct evolutionary path (Fig. S6). In contrast, the adjacent genes *hrpW1* and *avrE1* have an evolutionary history congruent with their respective phylogroups. The *Psa1-6 hopM1* alleles are all significantly different to the previously characterised *hopM1* in *Pto* DC3000, and this variation may be associated with the functional deletions in the *shcM* sequence.

### HopR1 and AvrE1 are additively required for *Psa3* virulence

Effectors outside the CEL locus have been reported to be functionally redundant with those in the CEL of *Pto* DC3000 (Cunnac et al., 2011; Kvitko et al., 2009). *Psa* alleles of *hopAA1-1* from the CEL and plasmid-borne *hopAA1-2*, though highly homologous to their counterparts in *Pto* DC3000, have both been pseudogenised by a 14 nucleotide or miniature inverted-repeat transposable element (MITE) insertion, respectively (Table 1; Fig. S7A) (Poulter et al., 2018; Templeton et al., 2015). *Psa3 hopR1*, however, appears to be fully functional and highly homologous (94.2% and 95.9% nucleotide and amino acid identities, respectively) to *hopR1* from *Pto* DC3000 (Fig. S7B). Interestingly, *hopR1* sequences within the *Psa* lineage appear to group into two clades, with the *Psa3* V13 allele grouping with those from *Pto* DC3000, biovars 2 and 6, as well as *Pfm* (Fig. S8). The more distantly related *hopR1* sequences from *Psa1* and *Psa5*, and closely related *Pmp*, appear to be significantly different in sequence, but is not characterised further here.

Since HopR1 was previously found to be redundantly required with CEL effectors in *Pto*, HopR1 function in *Psa3* V13 virulence was investigated. A *hopR1* knock-out in *Psa3* V13 (*ΔhopR1*), assessed by *in planta* growth, showed that *hopR1* is required for full *Psa3* virulence in ‘Hort16A’ leaf tissues at both 6 and 12 dpi (Fig. 7A). This was mirrored in disease symptomology with the *ΔhopR1* mutant displaying reduced lesions and plant death at 50 dpi (Fig. 7B). Intriguingly, a double CEL and *hopR1* knock-out strain (*ΔCEL/ΔhopR1*) was more severely impacted than *ΔCEL* alone, as assessed by both *in planta* growth as well as disease symptom development (Fig. 7C-D). This demonstrates additive contributions to *Psa3* virulence from both AvrE1 and HopR1, in a non-redundant manner.

**Figure 7.**
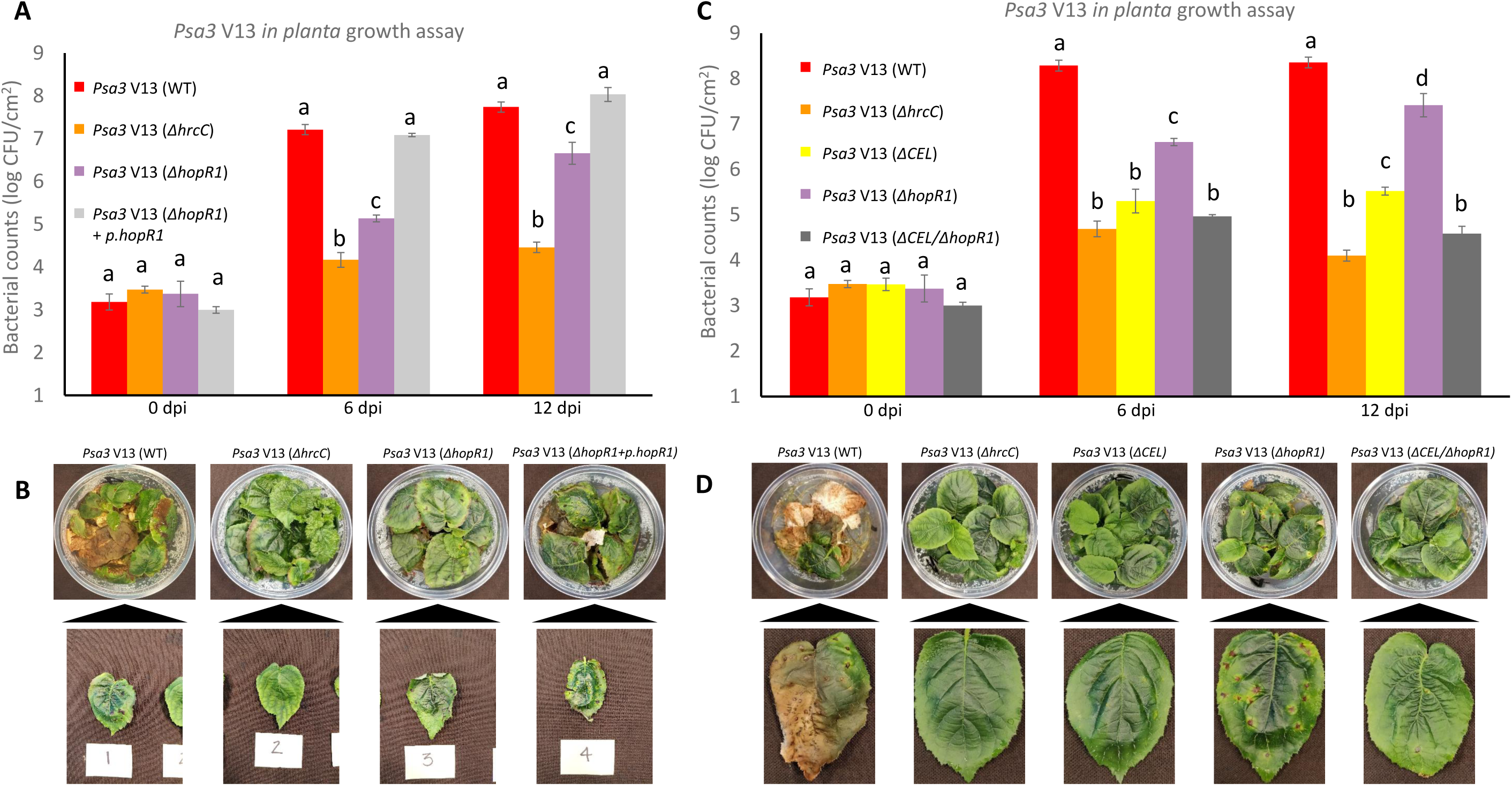
*hopR1* is non-redundantly required for virulence of *Pseudomonas syringae* pv. *actinidiae* ICMP 18884 biovar 3 (*Psa3*). **(A)** *Psa3* V13 (wildtype), *ΔhrcC* or *ΔhopR1* mutants, were inoculated by flooding at ∼10^6^ colony forming units; CFU/mL on *Actinidia chinensis* var. *chinensis* ‘Hort16A’, and bacterial growth was determined 0, 6, and 12 days post-inoculation (dpi). Error bars represent standard error of the mean from four pseudobiological replicates. Letters above the bars indicate results of a one-way analysis of variance (ANOVA) followed by Tukey’s HSD post-hoc test with samples per time point assigned different letters if significantly different (*P* < 0.05). The experiment was conducted three times with similar results. **(B)** Symptom development in ‘Hort16A’ at 50 dpi for strains inoculated in (A). Photographs taken of a single representative pottle and a representative leaf displaying typical disease lesion symptoms if present. **(C)** *Psa3* V13 (wildtype), *ΔhrcC, ΔCEL, ΔhopR1* mutants, or *ΔCEL/ΔhopR1* double mutant, were inoculated by flooding at ∼10^6^ CFU/mL on ‘Hort16A’, and bacterial growth was determined 0, 6, and 12 dpi. Error bars represent standard error of the mean from four pseudobiological replicates. Letters above the bars indicate results of a one-way ANOVA followed by Tukey’s HSD post-hoc test with samples per time point assigned different letters if significantly different (*P* < 0.05). The experiment was conducted three times with similar results. **(D)** Symptom development in ‘Hort16A’ at 50 dpi for strains inoculated in (C). Photographs taken of a single representative pottle and a representative leaf displaying typical disease lesion symptoms, if present.

## Discussion

Herein, *Psa3 in planta* growth (pathogen fitness) and symptom production (virulence) encoded by the evolutionary conserved CEL locus have been linked largely to effector AvrE1. The contribution to *in planta* growth and virulence is particularly dependent on the AvrE1 chaperone ShcE combination, since supplying plasmid-encoded AvrE1 without its chaperone only partially restored *in planta* growth and still lacked obvious symptom production long term. Interestingly, unlike a strong virulence contribution seen for HopM1 in *Pto* DC3000 (Badel et al., 2003, 2006), HopM1 in *Psa3*, with secretion rescued by ShcM_*Pto*_, had a small but significant reduction in pathogen fitness and no significant effect on virulence. Meanwhile, HopR1 was associated with a significant role in pathogen fitness, but only a minor role in virulence. Importantly, the fitness and virulence of *Psa3* V13 on susceptible kiwifruit ‘Hort16A’ has, for the first time, been shown to be dependent on functions of both HopR1 and AvrE1, both part of the same large family of virulence-associated effectors (Kvitko et al., 2009).

Effectors from the *P. syringae* species complex were recently collectively re-analysed to determine critical effectors that are associated with pathogenicity (Dillon et al., 2019a). Interestingly, of the four effectors considered part of the core genome of *P. syringae* (*avrE, hopB, hopM*, and *hopAA*), three are part of the CEL. Furthermore, only four effectors (*hopI, hopW, hopAH*, and *hopAS*) were more common than *hopR1* in *P. syringae* strains, indicating that CEL and associated effectors appear to be critical for this group of plant-pathogenic bacteria. In fact, Xin et al. (2018) suggest one of the earliest acquisitions of *P. syringae* on its evolutionary journey to pathogenicity would have been an AvrE1 and/or HopM1-like effector for water acquisition in the apoplast. Large-scale studies in *Arabidopsis* have identified that, of all environmental strains present on this model plant, the lineage of most successful bacterial colonisers (and potential pathogens) all carried a functional T3S and largely possessed *avrE1* (Karasov et al., 2018). Environmental isolates of *P. syringae* have been found to possess all three CEL effectors (*avrE1, hopM1*, and *hopAA1*) as well as *hopR1* in bulk-sequenced assemblies and both *avrE1* and *hopAA1* are conserved in over 95% of *P. syringae* species with *hopM1*, as well being present in a majority (Dillon et al., 2019b; Monteil et al., 2013).

In their large-scale environmental and agricultural sampling study, Dillon et al. (2019b) proposed that conserved ecologically and evolutionarily significant loci (like the CEL) are sites of increased inter-phylogroup recombination, amplifying genetic cohesion in the *P. syringae* species complex. The *Psa* lineage (a subset of phylogroup 1b defined here as strains including *Psa, Pfm*, and *Pmp*) alleles of *hopM1* appear to bear hallmarks of such a recombination event (Fig. S3) and may be driven by selection for evasion of host immunity in their native plant hosts. The variation in the *hopM1* sequence compared to flanking *avrE1* and *hrpW1* sequences suggests either a recombination event substituting the *hopM1* region specifically in the *Psa* lineage strains, or indicates evolutionary diversifying selection on the *hopM1* gene that has been noted previously for this gene across a number of *P. syringae* pathovars (Baltrus et al., 2011). Despite possessing an apparently full length amino acid sequence, *Psa3 hopM1* appeared to be non-functional due to a truncation in its chaperone *shcM*, unlike previously published non-functional *hopM1* alleles in the *Pto* lineage with mutations in the effector sequence (Cai et al., 2011). The loss of function suggests independent selection on *Psa* biovars during growth on kiwifruit (or possibly another shared host) for loss of HopM1 function. Nevertheless, when chaperone function was restored in *Psa3* V13, both in *cis* and in *trans*, the HopM1-dependency of reduced *in planta* growth was reminiscent of proposed HopM1-triggered recognition by tomato plants (Cai et al., 2011). This highlights that effector variation through evolutionary pressure encompasses both its native gene sequence as well as associated accessory genes required for efficient delivery into plant cells, a parameter underexplored in current research. Future evolutionary studies examining presence/absence of effectors should consider chaperone presence, both in *cis* (present in operon) and in *trans* (distal genomic presence), and sequence variation in the chaperone that could further impact on delivery efficiency, in the process of assessing an effector’s role in pathogenicity.

*P. syringae* pathogens that possess the canonical tripartite pathogenicity island have at least one of the conserved effectors *hopM1, avrE1*, or *hopAA1-1*, indicating a critical function for these effectors in pathogenicity (Alfano et al., 2000; DebRoy et al., 2004; Dillon et al., 2019a). Similar critical roles for CEL effector orthologs in multiple bacterial plant pathogens including *Erwinia* spp., *Pantoea* spp., *Pectobacterium* spp., and *Dickeya* spp. have been described (Alfano et al., 2000; Degrave et al., 2015). An *avrE1*-like effector was also recently found in the oomycete pathogen *Hyaloperonospora arabidopsidis* suggesting a conservation and functional role beyond bacterial pathogens (Deb et al., 2018). Effectors from the *Pto* DC3000 CEL show redundancy in their ability to promote disease, including *hopM1* and *avrE1* (Kvitko et al., 2009). Recently, further evidence of redundancy in the CEL was discovered through the fact that the same resistance protein that recognises AvrE1 in *Arabidopsis* also recognises HopAA1, suggesting a shared mechanism of action (e.g. a shared target monitored by this resistance gene) for these two effectors (Laflamme et al., 2020). Additionally, HopR1 in *Pto* DC3000 has been shown to be part of a complex of effectors in the same redundant effector group (REG) as AvrE1 and HopM1, albeit in a host-specific manner (Kvitko et al., 2009).

Intriguingly, *Psa3* AvrE1 and HopR1 appear to be functionally independent (Figure 7). In fact, the additive nature of AvrE1 and HopR1 contributions to *Psa3* pathogenicity indicate that unlike their orthologs in *Pto* DC3000, the *Psa3* versions do not appear to share a REG, at least for the susceptible kiwifruit host ‘Hort16A’ plants tested. This may suggest they have target(s) in kiwifruit that do not overlap. The remainder of CEL and associated effectors appear to be largely pseudogenised and non-functional in *Psa3*; HopM1 is nonfunctional due to truncation of its chaperone, while HopAA1-1 and HopAA1-2 are truncated through independent mutations. This suggests that AvrE1 alone contributes to virulence in the CEL-associated functions, and HopR1 independently contributes to an additional virulence function, making both attractive targets for durable resistance breeding.

While CEL effectors are required for pathogenicity, their precise role in disease establishment is still currently being elucidated. *Pto* DC3000 CEL (and associated) effectors HopM1 and AvrE1 have been found to suppress salicylic acid-mediated immunity by targeting signaling-related protein phosphatase 2A (DebRoy et al., 2004; Jin et al., 2016), hamper immunity-related secretion (Nomura et al., 2011), and induce water-soaking (Xin et al., 2016). HopAA1-1 is associated with formation of necrotic lesions and is a virulence factor sharing partial redundancy for disease symptom production with the phytotoxin coronatine biosynthetic gene *cmaL* (Munkvold et al., 2009; Worley et al., 2013). This is reminiscent of the symptom-associated role for AvrE1 and partial redundancy between *avrE1* and *hopAA1-1* seen earlier (Badel et al., 2006; Cunnac et al., 2011), and mirrored in shared recognition by the *CAR1* resistance gene in *Arabidopsis* (Laflamme et al., 2020). No conclusive function or mechanism of action for HopR1 has been described to date. However, HopR1 was recently found to be able to suppress HopQ1-triggered cell death in a strain-specific manner, suggesting an anti-immunity role (Zembek et al., 2018).

As yet, despite all key *Pto* DC3000 CEL and associated REG (CEL-REG) effectors triggering cell death in *N. benthamiana*, the resistance gene(s) recognising HopM1, HopR1, HopAA1 and/or AvrE1 is unknown (Wei et al., 2007, 2018). Nevertheless, the CEL effectors’ ability to render *Pto* DC3000 avirulent on *N. benthamiana* is hampered by the presence of suppressor effectors HopI1 and AvrPtoB (Wei et al., 2018). This suggests that while CEL-REG effectors may be conserved for function in a number of different plant pathogens, a required suppressor-REG also exists in these bacterial pathogens to counter the plant hosts’ ability to recognise and trigger immunity against the CEL-REG effectors. This may also be true for *Psa* and is a significant focus of current research. The interest in this proposed suppressor-REG is largely because functionally redundant suppressor effectors make attractive targets for future programs for plant resistance breeding, since resistance to these suppressor effectors, in turn, is likely to be durable due to their requirement in masking immunity triggered by multiple virulence-critical CEL and associated effectors.

Understanding pathogen emergence in new crops such as kiwifruit requires the discrimination of non-redundant disease-critical elements from those that may be redundant. Most *P. syringae* strains appear to have a narrow spectrum of hosts they may infect (Morris et al., 2019). Interestingly, kiwifruit appears highly resistant to most characterised *P. syringae* pathovars, with the obvious exception of *Psa* strains that appear to be pathogenic on a number of plants, suggesting that they possess an intrinsic ability to cause more virulent disease (Morris et al., 2019). Thus, understanding the core effectors in *Psa* that are required for pathogenicity and associated with increased virulence as well as the redundancy in this system is likely to impact on our understanding of other *P. syringae* pathogens that infect and trigger virulent disease in a large number of plant species.

## Experimental Procedures

### Bioinformatics and sequences

Genome sequences for *Psa3* ICMP 18884 (*Psa3* V13) and other strains in this study were obtained from NCBI Genbank (CP011972.2 and CP011973.1). The *Psa3* V13 genome was annotated previously (Templeton et al., 2015). Sequences for T3Es from *Psa3* V13, other *Psa* lineage strains, and type strains from phylogroups 1-3 were analysed on Geneious™ R11 software (https://www.geneious.com; Biomatters, NZ) with built-in Geneious DNA and amino acid sequence alignments, tree building (RAxML v8 with 100 bootstrapping replicates), and annotation tools.

### Bacterial strains and growth conditions

Bacterial strains and plasmids used in this study are listed in Supplementary Table S1. *Psa3* V13 strains were grown in Lysogeny Broth (LB) at 20 °C with shaking at 200 rpm. *Escherichia coli* strains were grown in LB with appropriate antibiotics at 37°C. The concentrations of antibiotics used in selective media were: kanamycin, 50 μg/mL; gentamicin, 25 μg/mL; nitrofurantoin, 12.5 μg/mL; cephalexin, 40 μg/mL; chloramphenicol, 10 μg/mL; and tetracycline, 5 μg/mL (all from Sigma-Aldrich). Plasmids were transformed into electrocompetent *Psa3* (Choi et al., 2006; Mesarich et al., 2017) or *E. coli* by electroporation using a BioRad Gene Pulser Xcell™ and recovered for 1 hour in LB before plating on selective media.

### Effector gene knock-out

To make the CEL or *hopR1* deletions in *Psa*3 V13, an approach derived from that described for *Pto* DC3000 was utilised (Kvitko et al., 2009). Briefly, DNA fragments containing the upstream (∼1 kb) and downstream (∼1 kb) regions of the effector (T3E) of interest were PCR-amplified using primer pairs Psa_(T3E)-KO_UP-F/ Psa_(T3E)-KO_UP-R and Psa_(T3E)-KO_DN-F/Psa_(T3E)-KO_DN-R, respectively (Supplementary Table S2), with the UP-R and DN-F primers carrying *Xba*I restriction enzyme sites. Each PCR fragment was then gel-purified using an EZNA gel extraction kit (Custom Science, NZ) and then digested with *Xba*I, re-purified with an EZNA PCR product purification kit and then ligated to form the ∼2 kb KO fragment. The ∼2 kb fragment containing both the upstream and downstream fragments for each T3E was then re-amplified by PCR using primers Psa_(T3E)-KO_UP-F and Psa_(T3E)-KO_DN-R. This PCR product was cloned into the *Eco53k*I blunt-end restriction enzyme site of pK18mobsacB (Schäfer et al., 1994) that had first been mutagenised to remove the non-MCS *Eco53k*I site to generate pK18B-E. The pK18B-E vector carrying the ∼2 kb KO fragment, called pΔ(T3E), was transformed into *E. coli* DH5α, plated on X-gal/IPTG/kanamaycin LB agar plates for blue/white selection and positive transformants confirmed by Sanger sequencing (Macrogen, South Korea). For each T3E knock-out, *Psa3* V13 was transformed with the relevant pΔ(T3E) construct and transconjugants were selected on LB plates with nitrofurantoin, cephalexin and kanamycin. Selected colonies were subsequently streaked onto LB plates containing 10% (w/v) sucrose to counter-select plasmid integration. *ΔCEL* or *ΔhopR1* mutants were screened using colony PCR with primers Psa_(T3E)-KO_Check-F and Psa_(T3E)-KO_Check-R, and sent for Sanger sequencing with the cloning Psa_(T3E)-KO_UP-F and Psa_(T3E)-KO_DN-R primers, as well as further confirmed using internal T3E-specific primers and plating on kanamycin-containing medium to confirm loss of the *sacB* gene.

### Effector gene knock-in

The sequences for *hrpW1, hopM1* (with chapeone *shcM*), and *avrE1* (with chaperone *shcE*) were synthesised with their native HrpL box promoter sequences, synonymous mutations to remove internal *Mlu*I sites, and flanking *Asc*I restriction enzyme sites added in the pUC57 vector (GenScript, USA). These pUC57 vectors carrying the knock-in gene of interest were then single-pot cloned (with *Asc*I, *Mlu*I, and T4 DNA ligase enzymes in T4 buffer; NEB) into the *Mlu*I site in the pΔCEL vector. The pΔCEL vector carrying the knock-in gene, called pΔCEL+hrpW1, pΔCEL+shcM:hopM1, or pΔCEL+shcE:avrE1, was transformed into *E. coli* DH5α, plated on kanamaycin LB agar plates and screened by colony PCR for the inserted gene. Positive transformants were confirmed by Sanger sequencing (Macrogen, South Korea). For each gene knock-in, *Psa3* V13 *ΔCEL* was transformed with the relevant p*ΔCEL*+(gene-of-interest) construct and transconjugants were selected on LB agar plates with nitrofurantoin, cephalexin and kanamycin. Selected colonies were subsequently streaked onto LB agar containing 10% (w/v) sucrose to counter-select plasmid integration. Insertion mutants, *ΔCEL+hrpW1, ΔCEL+hopM1*, or *ΔCEL+avrE1*, were screened using colony PCR with primers Psa_(T3E)-KI_Check-F and Psa_(T3E)-KI_Check-R, and sent for Sanger sequencing with internal gene-specific primers, as well as further confirmed by plating on kanamycin-containing medium to confirm loss of the *sacB* gene.

### *shcM*_*Pto*_ complementation

*shcM*_*Pto*_, including its HrpL box promoter from *Pto* DC3000, was PCR-amplified using shcM_Pto-F and shcM_Pto-R primers (for *trans*-complementation) or shcM_Pto-GG-F and shcM_Pto-GG-R golden gate primers (for *cis*-complementation by assembly into the modified broad host-range vector pBBR1MCS-5-GG:*avrRsp4*_*pro*_ (Jayaraman et al., 2017)) (Supplementary Table S2). The resulting PCR fragment was gel-purified as above. For the *trans*-complementation construct, the extracted PCR product was blunt-end-ligated into the *Eco53k*I site of broad host-range vector pBBR1MCS-5 (Kovach et al., 1995). For the *cis*-complementation construct, the extracted PCR product was blunt-end-ligated into the *Eco53k*I site of shuttle vector pICH41021 (Engler et al., 2008; Weber et al., 2011). Both constructs were transformed into *E. coli* DH5α, plated on X-gal/IPTG-containing (for blue/white selection) LB agar plates with gentamicin (*trans*) or ampicillin (*cis*) selection, and positive transformants confirmed by Sanger sequencing (Macrogen, South Korea). The pICH41021-*shcM*_*Pto*_ construct was then used to golden gate-assemble the pBBR1MCS-5:*avrRsp4*_*pro*_:*shcM*_*Pto*_:*hopM1* construct with a 6xHA tag added to C-terminus of HopM1, as done previously for other effector constructs (Choi et al., 2017; Jayaraman et al., 2017). Both *trans*- and *cis*-complementation constructs were transformed into relevant *Psa3* strains by electroporation, as described previously and transformants screened for presence of *shcM*_*Pto*_ by gene-specific colony PCR, followed by digest by *Nde*I (specific for *Psa3* allele of *shcM*).

### Infection assays

*Psa* infection assays were based on previous conditions (McAtee et al., 2018). *Actinidia chinensis* var. *chinensis* ‘Hort16A’ plantlets, grown from axillary buds on Murashige and Skoog rooting medium without antibiotics in sterile 400-mL plastic tubs (pottles), were purchased from Multiflora (Auckland, New Zealand). Plantlets were grown at 20 °C under Gro-Lux fluorescent lights under long-day conditions (16 h:8 h; light:dark) and used when the plantlets were between 8-12 weeks old. Overnight cultures of wildtype or mutant strains of *Psa3*were pelleted at 6000 *g*, re-suspended in 10 mM MgSO_4_, cell density determined by measuring the absorbance at 600 nm, and reconstituted at Abs_600_ = 0.05 (∼10^7^ colony forming units; CFU/mL, determined by plating) in 500mL of 10 mM MgSO_4_. Surfactant Silwet™ L-77 (Lehle Seeds, Round Rock, TX, USA) was added to the inoculum at 0.0025% (v/v) to facilitate leaf wetting. Pottles of ‘Hort16A’ plantlets were flooded with the inoculum, submerging the plantlets for three minutes, drained, sealed, and then incubated under previously described plant growth conditions.

### Pathogen growth and symptom development

*Psa3* V13 *in planta* growth assays were based on previous conditions (McAtee et al., 2018). Briefly, leaf samples of four leaf discs per pseudobiological replicate, taken randomly with a 1-cm diameter cork-borer from three plants, were harvested at 2 h (day 0), day 6, and day 12 post-inoculation. All four replicates per treatment, per timepoint were taken from the same pottle. To estimate *Psa3* growth inside the plant, the leaf discs were surface-sterilised, placed in Eppendorf tubes containing three sterile stainless-steel ball bearings and 350 μL 10 mM MgSO_4_, and macerated in a Storm 24 Bullet Blender (Next Advance, NY, USA) for two bursts of 1 min each at maximum speed. A ten-fold dilution series of the leaf homogenates was made in sterile 10 mM MgSO_4_ until a dilution of 10^− 8^ and plated as 10 μL droplets on LB medium supplemented with nitrofurantoin and cephalexin. After 2 days of incubation at 20°C, the CFU per cm^2^ of leaf area was ascertained from dilutions. Plasmid loss was investigated by plating the pBBR1MCS-carrying strains on LB agar medium with and without gentamicin (Gm or non-Gm). To observe pathogenic symptoms on the plants, infected pottles were kept up to 50 days post-inoculation and photographs taken of pottles and a representative infected leaf. Infection severity was qualitatively assessed based on typical symptoms: necrotic leaf spots, chlorotic haloes, leaf death, and plant death. Each of these growth assay experiments were conducted at least three times.

### *In planta* effector secretion assay

Broad host-range plasmid constructs (pBBR1MCS-5-GG:*avrRps4*_*pro*_) of each *Psa3* T3E were transformed by electroporation into *P. fluorescens* Pf0-1(wildtype) or *P. fluorescens* Pf0-1(T3S) strain (Thomas et al., 2009) and plated on selective medium with chloramphenicol, gentamicin, and tetracycline (only +T3S strains). Positive transformants were confirmed by gene-specific colony PCR. Pf0-1(T3S) carrying empty vector or *Psa3*-T3E constructs, were streaked from glycerol stocks onto LB agar plates with antibiotic selection and grown for 2 days at 28°C. Bacteria were then harvested from plates, resuspended in 10 mM MgSO_4_ and diluted to required Abs_600_ = 1.5 (∼10^9^ CFU/mL). Infiltrations were carried out on fully expanded leaves of 4- to 5-week-old *N. benthamiana* with a blunt-end syringe, on two half-leaf sections of *N. benthamiana* leaves (replicates). Leaf samples were harvested at 6 h post-infiltration and snap frozen in liquid nitrogen, ground with mortar and pestle, boiled in 1xLeamli buffer with DTT and run on SDS-PAGE for immunoblot for presence of the 6xHA-tagged T3E using α–HA antibody. Membranes were subsequently stained with Coomassie Brilliant Blue to visualise total protein for gel loading control.

### Defense gene expression by qPCR

Total RNA was extracted from four leaf discs per replicate (three replicates, each sampled from two independent flood-inoculated ‘Hort16A’ plantlets) via a Spectrum™ Plant Total RNA kit (Sigma Aldrich, NZ). RNA was DNase I (Sigma Aldrich, NZ) treated and cDNA was synthesised from RNA using the High-Capacity cDNA Reverse Transcription kit following manufacturer’s instructions (Thermo Fisher, NZ). Quantitative PCR (qPCR) was carried out on an Illumina Eco™ Real-Time PCR machine using the EvaGreen SsoFast™ qPCR mix (BioRad, NZ). Primers used for qPCR are listed in Supplementary Table S2.

## Acknowledgements

This work was funded (including a post-doctoral fellowship to JJ) by the Bio-protection Research Centre (Tertiary Education Commission). We would like to thank Jo Bowen (PFR), Erik Rikkerink (PFR), and Carl Mesarich (Massey University) for critically reading the manuscript.

## Supplementary methods

### *Nicotiana benthamiana* HR assays

Broad host-range plasmid constructs (pBBR1MCS-5) of each *Pseudomonas syringae* pv. *actinidiae* ICMP 18884 biovar 3 (*Psa3*) V13 T3E were transformed by electroporation into *P. fluorescens* Pf0-1 (wildtype) or *P. fluorescens* Pf0-1 (T3S) strain (Thomas et al., 2009) and plated on selective media with chloramphenicol, gentamicin, and tetracycline (only +T3S strains). Positive transformants were confirmed by gene-specific colony PCR. Pf0-1(T3S) carrying empty vector or *Psa3*-T3E constructs, were streaked from glycerol stocks onto LB agar plates with antibiotic selection and grown for 2 days at 28°C. Bacteria were then harvested from plates, resuspended in 10 mM MgSO_4_ and diluted to required Abs_600_ = 0.6 or 1 (∼10^8-9^ colony forming units; CFU/mL). Infiltrations were carried out on fully expanded leaves of 4- to 5-week-old *Nicotiana benthamiana* using a blunt-end syringe, on 2–3 leaves (replicates), with the full experiment repeated three times. Hypersensitive cell death response was assayed visually and photographs taken at 3 days post-infiltration (dpi).

### *In vitro* effector secretion assay

For detection of T3E secretion *in vitro*, protocols utilised are based on those described previously (Huynh et al., 1989; Roine et al., 1997). Briefly, *Psa3* V13 conserved effector locus mutant (*ΔCEL*) or *Psa3* V13 type III secretion system mutant (*ΔhrcC*) (Colombi et al., 2017) carrying the relevant broad-host range plasmid constructs (pBBR1MCS-5 or pBBR1MCS-2) were grown in rich LB medium with antibiotic selection overnight, pelleted at 6000 *g*, washed with *hrp*-inducing minimal medium supplemented with 10mM fructose (Huynh et al., 1989) and then re-suspended in *hrp*-inducing minimal medium and incubated for 6 h with shaking for *hrp* induction. Following *hrp* induction, cells were pelleted, supernatant carefully separated and passed through a 0.2 μm filter, and proteins from both pellet and supernatant were resolved by SDS-PAGE and immunoblotted for the presence of the 6xHA-tagged T3E using α–HA antibody.

**Figure S1.**
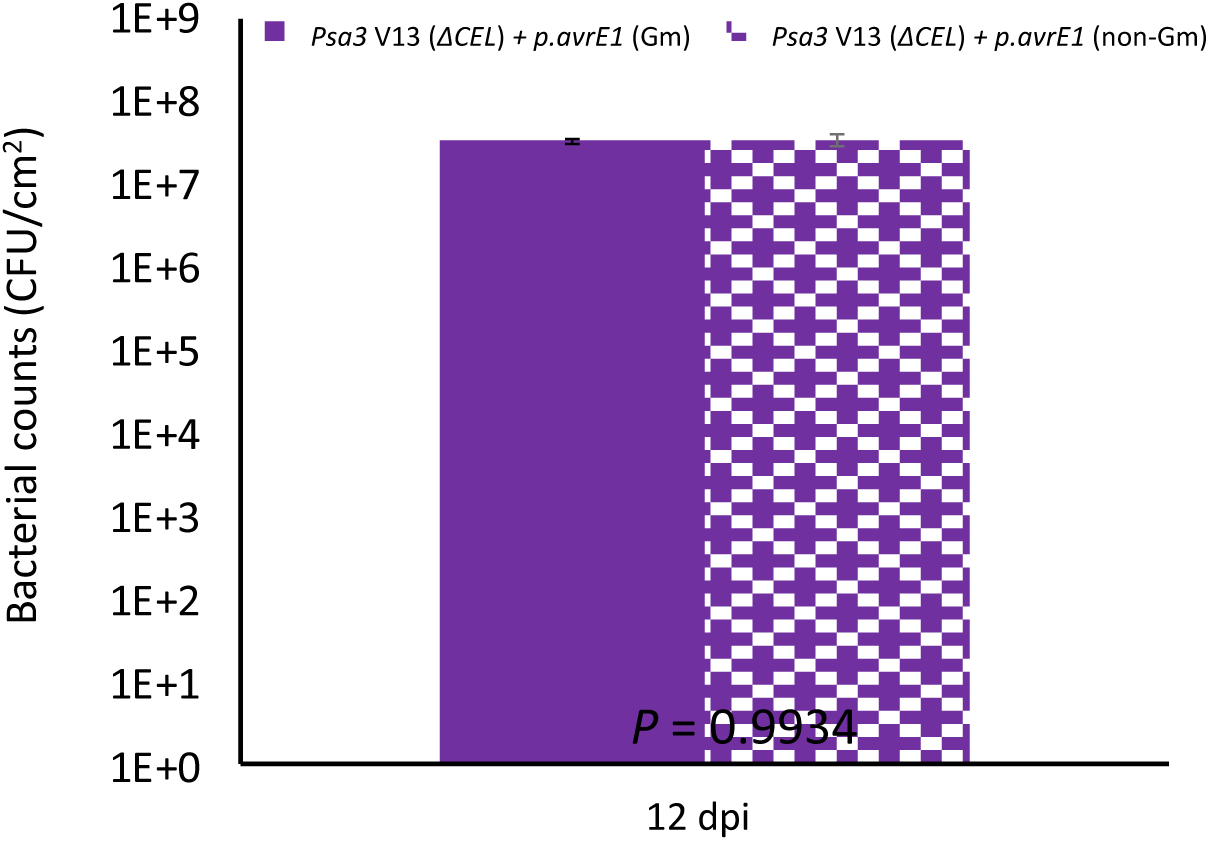
*avrE1* partial complementation of virulence of *Pseudomonas syringae* pv. *actinidiae* ICMP 18884 biovar 3 conserved effector locus mutant (*Psa3 ΔCEL*) is not due to plasmid loss. *Psa3 ΔCEL* carrying plasmid-borne *avrE1* infected *Actinidia chinensis* var. *chinensis ‘*Hort16A’ sampled 12 days post-inoculation were plated with and without gentamicin selection (Gm and non-Gm, respectively). Error bars represent standard error of the mean from four pseudobiological replicates. *P* value from a two-tailed Student’s *t*-test comparison between Gm and non-Gm are shown.

**Figure S2.**
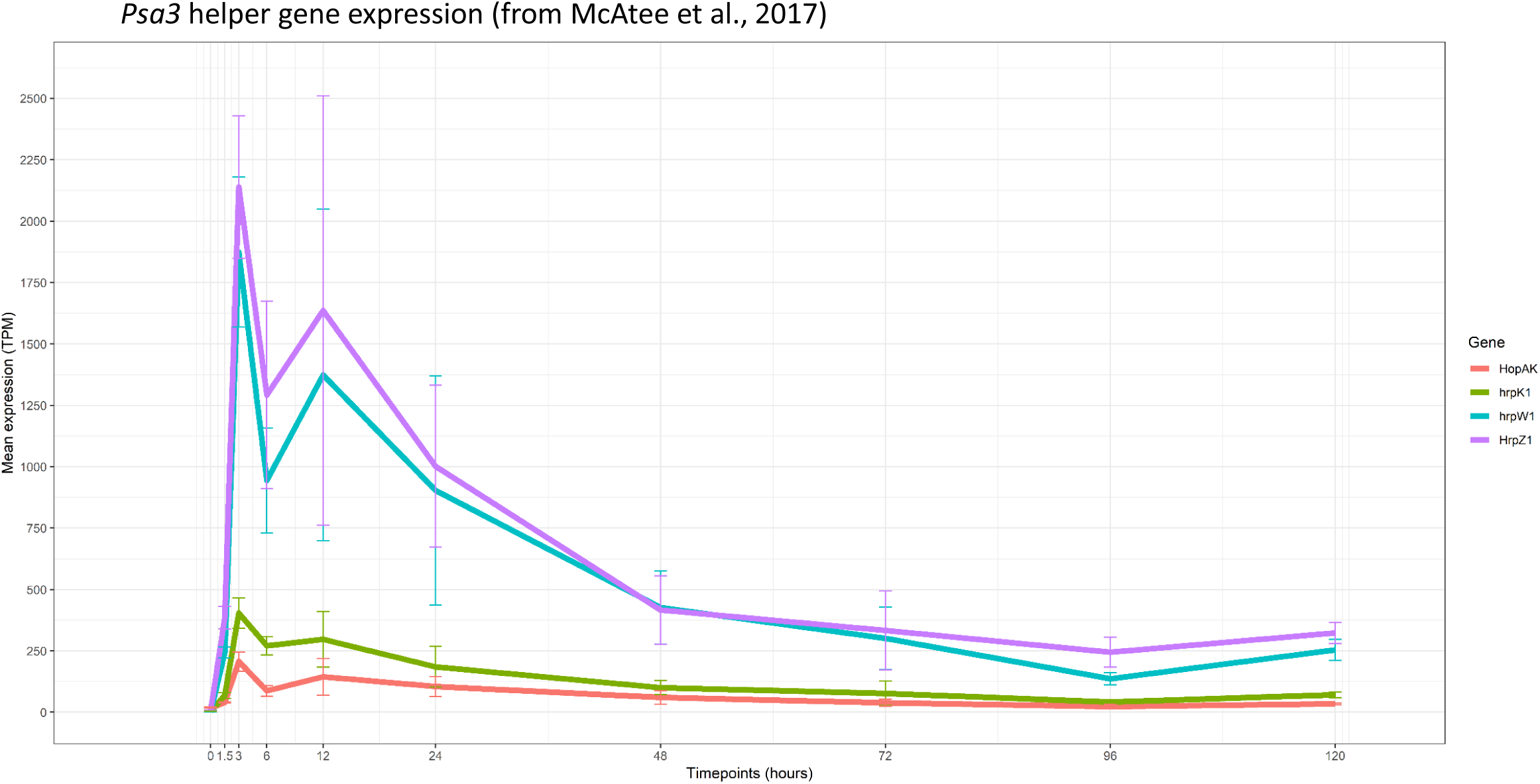
*hrp* (helper) gene expression during the first 5 days of infection. *Pseudomonas syringae* pv. *actinidiae* ICMP 18884 biovar 3 (*Psa3*) V13 infected on *Actinidia chinensis* var. *chinensis* ‘Hort16A’ plants were sampled at indicated times and assessed for bacterial gene expression by RNA-seq of four *hrp* helper genes *hopAK, hrpK1, hrpZ1*, and *hrpW1* located in the *hrp*/*hrc* cluster or the conserved effector locus (CEL) - *hrpW1*. Data is extracted from previously published study (McAtee et al., 2018). TPM; transcripts per million.

**Figure S3.**
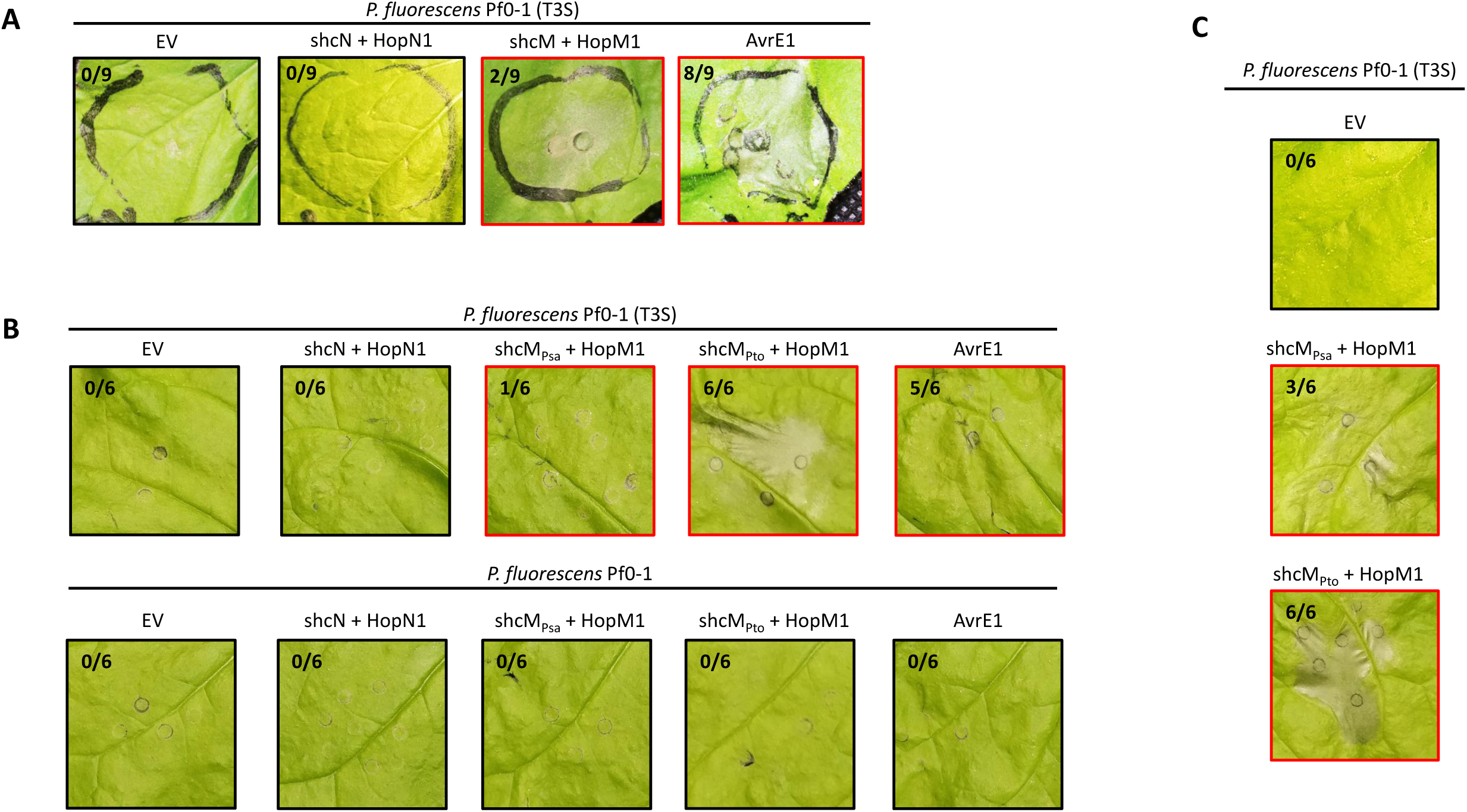
*shcM*_*Pto*_ facilitates HopM1 to trigger cell death in *Nicotiana benthamiana*. **(A)** *Pseudomonas fluorescens* (*Pfo*) Pf0-1 (T3S) carrying broad host-range vector pBBR1MCS-5 : *avrRps4* promoter : (Effector_*Psa*_) : 6xHA constructs (Jayaraman et al., 2017) for the effectors *avrE1* (without *shcE*), *hopM1* (with *shcM*), or *hopN1* (with *shcN*), or empty vector (EV) were infiltrated into *Nicotiana benthamiana* leaves at Abs_600_ = 1. **(B)** *Pfo* Pf0-1(T3S) or *Pfo* Pf0-1 (WT) carrying broad host-range vector constructs from (A) for empty vector (EV), *hopN1* (with *shcN*), *hopM1* (with chaperone *shcM*_*Psa*_ or *shcM*_*Pto*_), or *avrE1* (without *shcE*) were infiltrated into *N. benthamiana leaves* at Abs_600_ = 0.6. **(C)** *P. fluorescens* Pf0-1 (T3S) carrying pBBR1MCS-5 *hopM1* construct from (B) with *shcM*_*Psa*_ or *shcM*_*Pto*_ were infiltrated into *N. benthamiana leaves* at Abs_600_ = 1. All panels shown are representative of the symptoms from three independent experiments (counts indicated at top right of each panel) and are framed in red where any cell death was visible. All photographs were taken at 3 days post-infiltration.

**Figure S4.**
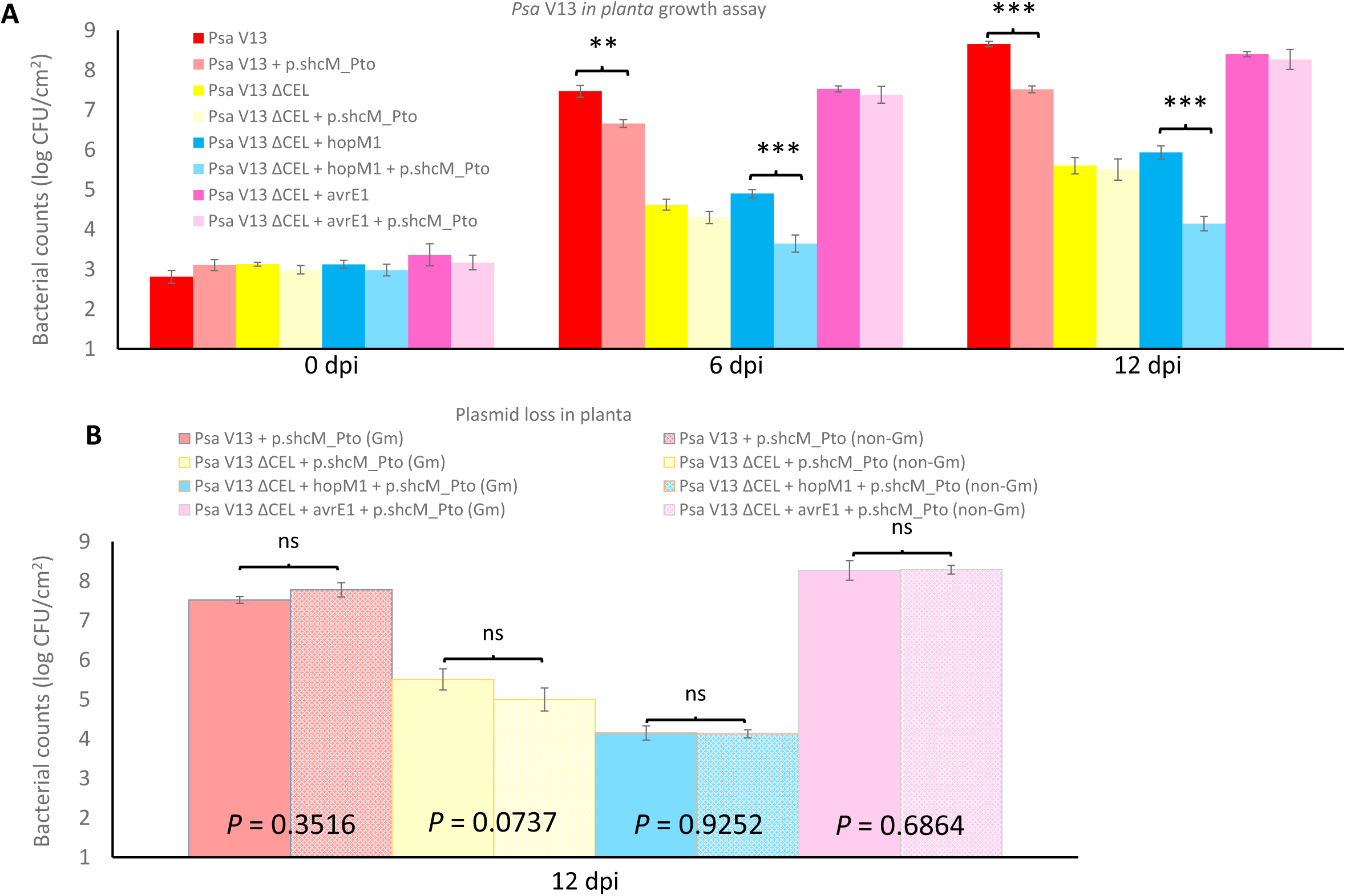
*Trans*-complementation with *shcM* from *Pseudomonas syringae* pv. *tomato* (*Pto*) DC3000 reduces virulence of *Pseudomonas syringae* pv. *actinidiae* ICMP 18884 biovar 3 (*Psa3*) V13 in a *hopM1*-dependent manner. **(A)** *Psa3* V13 (wildtype), *ΔCEL* mutant, *ΔCEL* with genomic knock-in *shcM:hopM1*, or *ΔCEL* with genomic knock-in *shcE:avrE1* with or without plasmid-borne *shcM*_*Pto*_, were flood-inoculated at ∼10^6^ colony forming units; CFU/mL into *Actinidia chinensis* var. *chinensis* ‘Hort16A’, and bacterial growth was determined 0, 6, and 12 days post-inoculation (dpi). Error bars represent standard error of the mean from four pseudobiological replicates. Asterisks indicate results of a two-tailed Student’s *t*-test between selected samples indicated; **(*P* < 0.01), ***(*P* < 0.001). The experiment was conducted two times with similar results. **(B)** *Psa3* V13-infected *‘*Hort16A’ leaf samples 12 dpi from (A) were plated with and without gentamicin selection (Gm and non-Gm, respectively). Error bars represent standard error of the mean from four pseudobiological replicates. *P* value from a two-tailed Student’s *t*-test comparison between Gm and non-Gm are shown.

**Figure S5.**
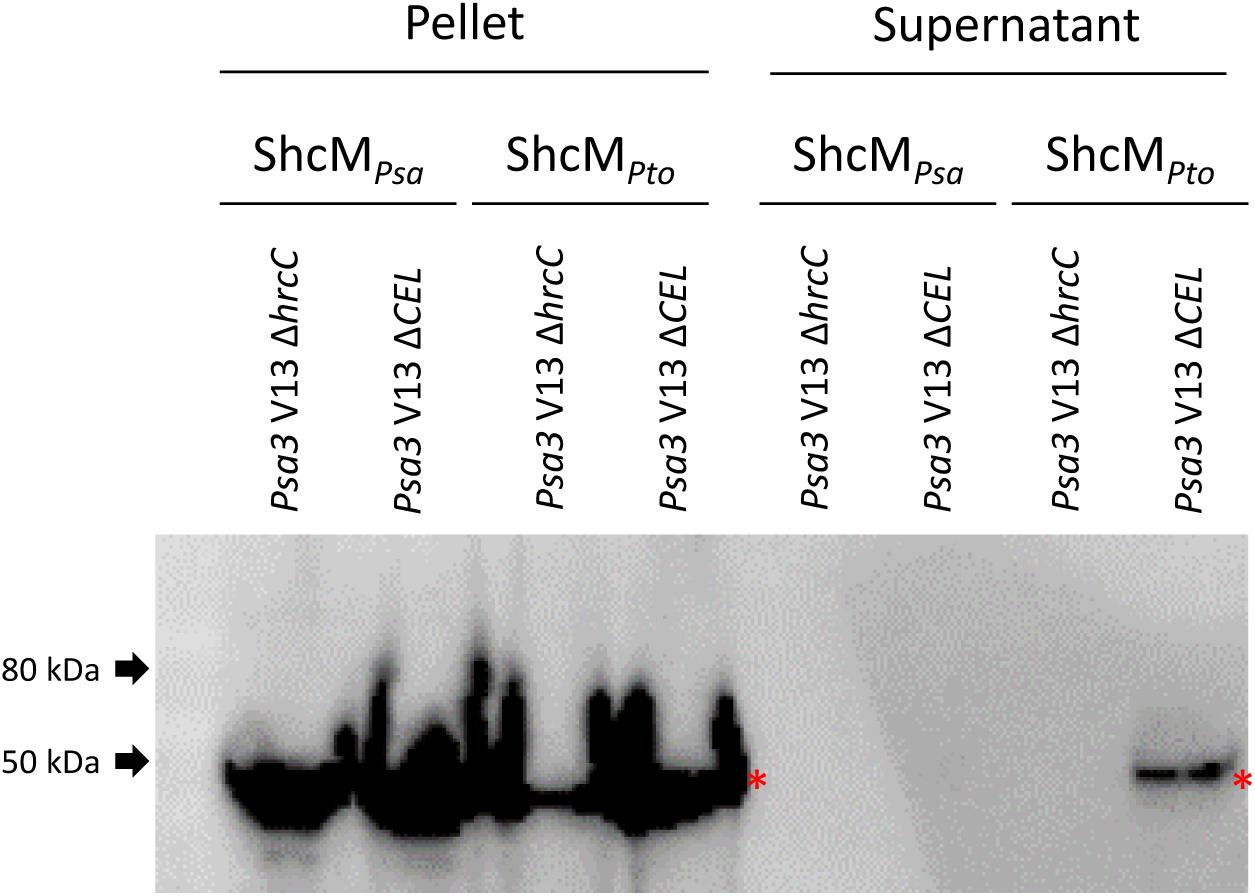
*Pseudomonas syringae* pv. *actinidiae* ICMP 18884 biovar 3 (*Psa3*) V13 carrying *shcM* from *Pseudomonas syringae* pv. *tomato* (*Pto*) DC3000 can deliver HopM1. *In vitro* secretion assay for HopM1 protein tagged with 6xHA from *Psa3 ΔCEL* or *Psa3 ΔhrcC* with *shcM*_*Psa*_ or *shcM*_*Pto*_. *Psa3* strains were inoculated at Abs_600_ = 0.5 and grown for 6 h on *hrp*-inducing minimal medium with fructose (Huynh et al., 1989). Equal volumes of pellet and supernatant for each strain were separated by centrifugation, boiled in 1xLeamli buffer and run on SDS-PAGE for immunoblot using α–HA antibody. The red asterisks indicate expected sizes for the tagged protein bands.

**Figure S6.**
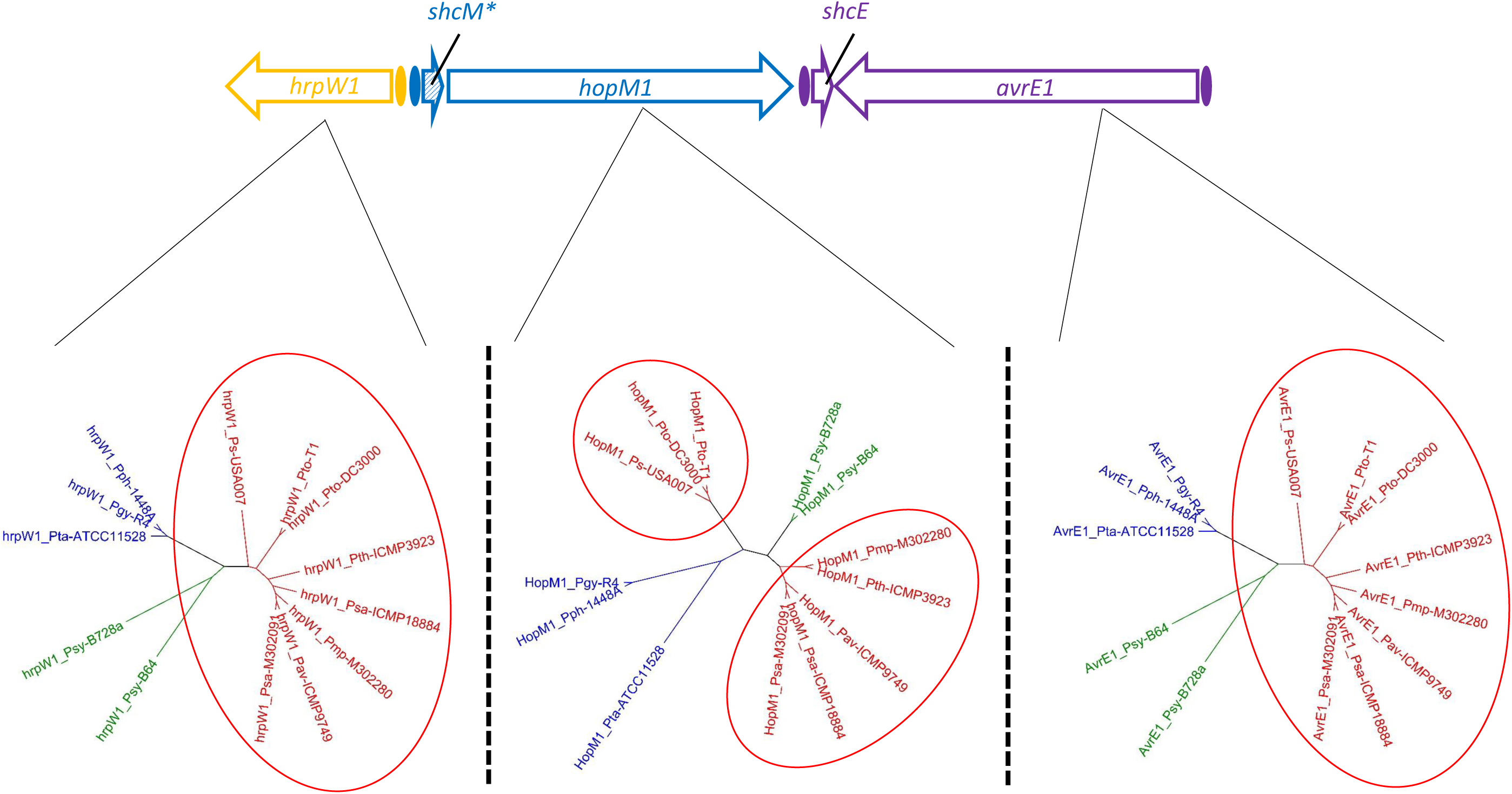
The *hopM1* lineage in *Pseudomonas syringae* phylogroup 1 is split into two lineages, unlike other conserved effector locus (CEL) genes around it. A maximum likelihood (RAxML v8, 100 bootstrap replicates) tree for *hrpW1, hopM1* (*shcM* indicated with an asterisk and hatched shading due to truncation in Bv3 allele), or *avrE1* gene sequences for representative members from the major plant pathogenic phylogroups of *P. syringae* compared to *P. syringae* pv. *actinidiae* ICMP 18884 biovar 3 (*Psa3*) V13 (from phylogroup 1). Phylogroup 1 (red): to *P. syringae* pv. *tomato* (*Pto*) DC3000, *Pto* T1, to *P. syringae* pv. *morsprunorum* (*Pmp*; a.k.a. *Pam*/*Psm* R2) MAFF 302280, *Psa* MAFF 302091. Phylogroup 2 (green): *P. syringae* pv. *syringae* (*Psy*) B64, and *Psy* B728A. Phylogroup 3 (blue): *P. syringae* pv. *tabacum* (*Pta*) ATCC 11528, *P. syringae* pv. *glycinea* (*Pgy*) R4, and *P. syringae* pv. *phaseolicola* (*Pph*) 1448A. Phylogroup 5 (pink): *P. syringae* pv. *maculicola* (*Pma*) ES4326.

**Figure S7.**
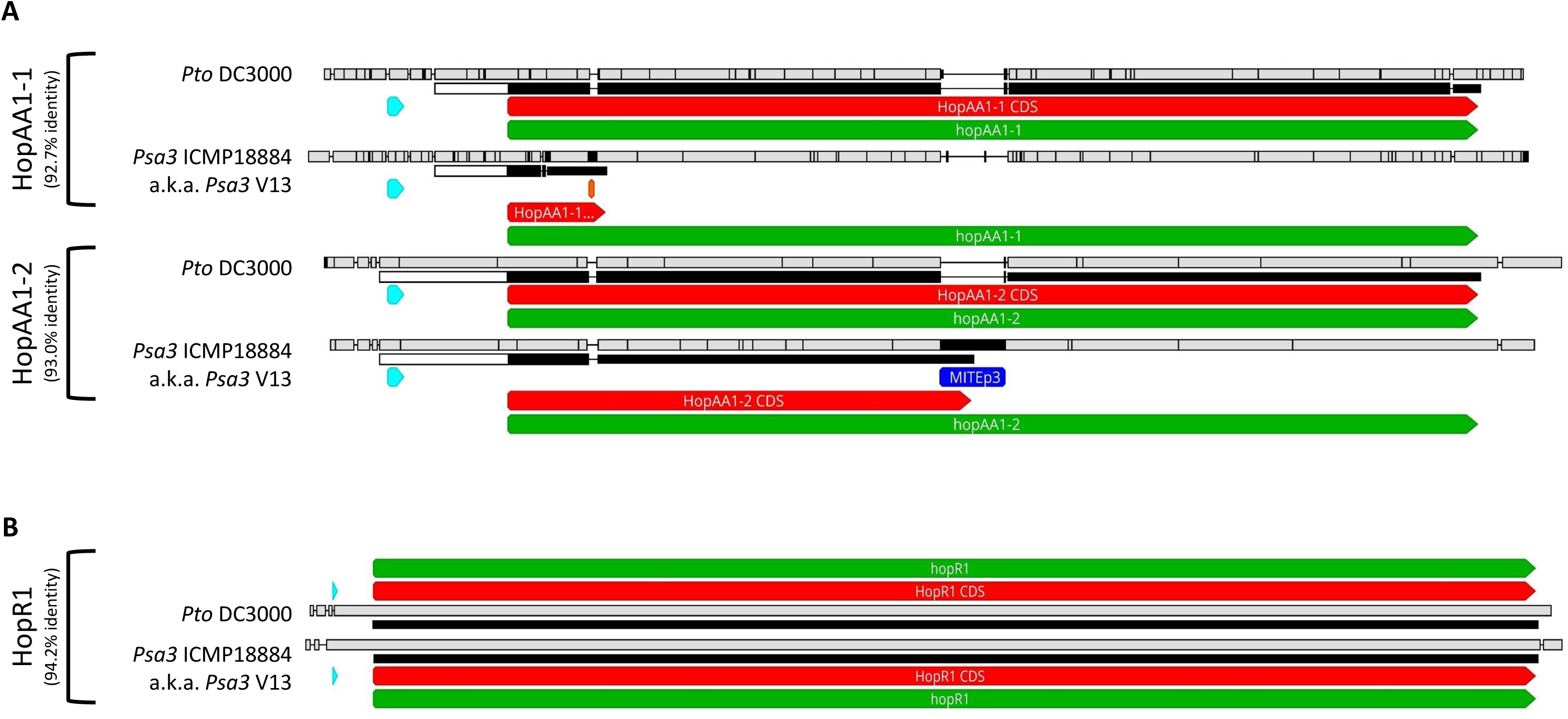
HopAA1 and HopR1 alleles in *Pseudomonas syringae* pv. *actinidiae* ICMP 18884 biovar 3 (*Psa3*). Nucleotide sequence alignment of the *hopAA1-1* and *hopAA1-2* regions **(A)** and *hopR1* **(B)** of *P. syringae* pv. *tomato* (*Pto*) DC3000 (*hopAA1-1*_*Pto*DC3000_ and *hopR1*_*Pto*DC3000_, respectively, set as reference) and *Psa3* ICMP 18884 (*Psa3* V13) as viewed in Geneious software. The top bar (black/white) in each panel represents nucleotide sequence similarity with the bar underneath representing amino acid similarity, red arrows indicate coding sequences, and green arrows indicate genomic regions (ORFs). The HrpL box promoters are represented by the blue arrowhead for each sequence.

**Figure S8.**
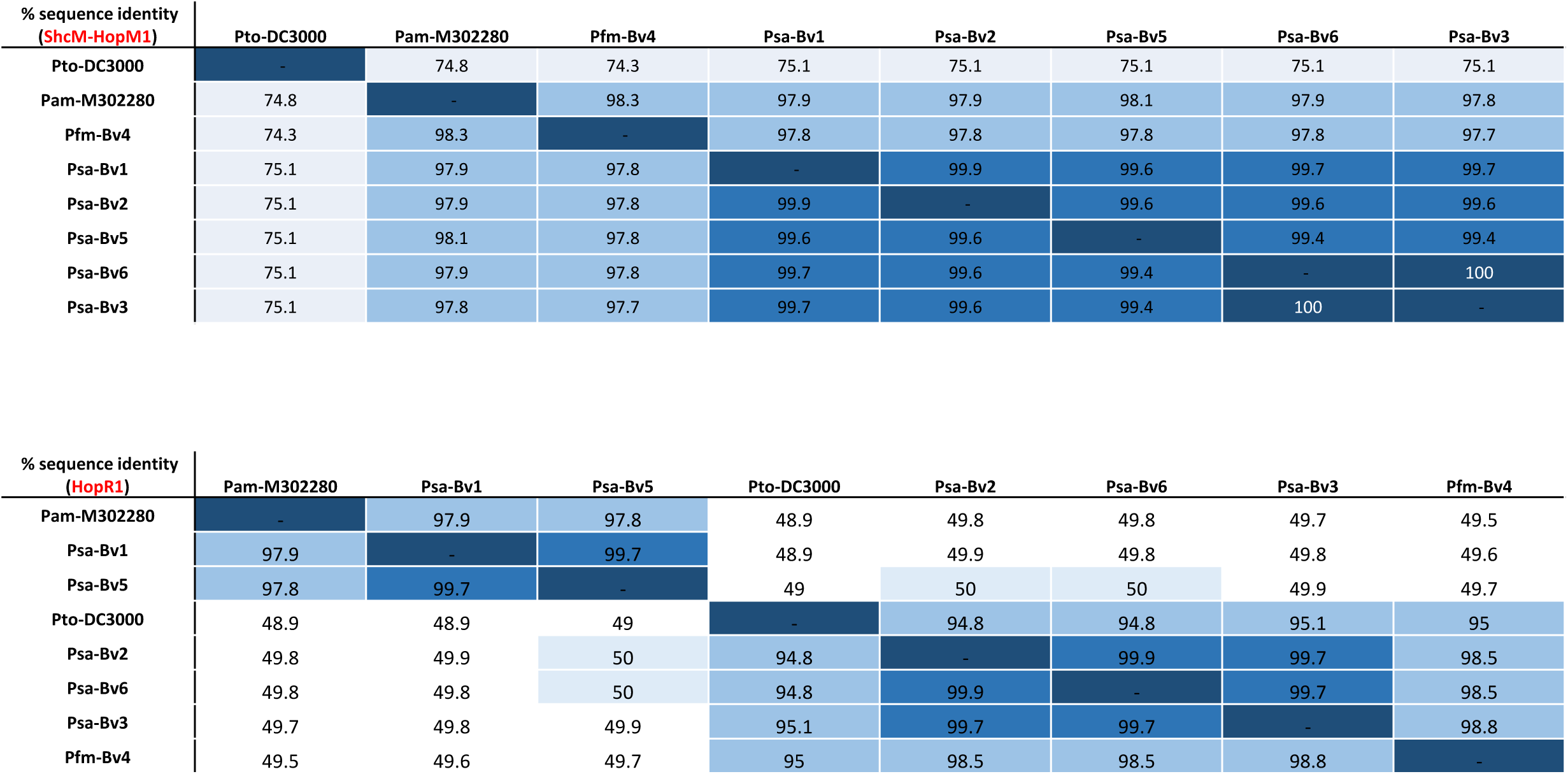
Alleles of HopM1 from the *Pseudomonas syringae* pv. *actinidiae* (*Psa*) lineage are closely related, while HopR1 alleles are broadly divided into two groups. Amino acid sequence identity comparison of HopM1 **(A)** or HopR1 **(B)** between *P. syringae* pv. *tomato* (*Pto*) DC3000, *P. syringae* pv. *morsprunorum* (*Pam*) MAFF 302280, *P. syringae* pv. *actinidifoliorum* (*Pfm*) ICMP 18803 (previously *Psa4*), *Psa1* ICMP 9855, *Psa2* ICMP 19073, *Psa5* MAFF 212063, *Psa6* MAFF 212141, and *Psa3* IMCP 18884.

**Supplementary Table S1.**

| Strain | Description | Reference |
| --- | --- | --- |
| *Psa3* V13 | *P. syringae* pv. *actinidiae* biovar 3 isolate NZ V-13 (*Psa3* V13; wildtype) | McCann et al., 2013 |
| *Psa3* V13 $\Delta hrcC$ | *Psa3* V13 *hrcC* knockout | Colombi et al., 2017 |
| *Psa3* V13 $\Delta CEL$ | *Psa3* V13 *CEL* knockout | This study |
| *Psa3* V13 $\Delta CEL$ + p.EV | *Psa3* V13 *CEL* knockout with empty vector | This study |
| *Psa3* V13 $\Delta CEL$ + p.*hopN1* | *Psa3* V13 *CEL* knockout with plasmid-borne *shcN* and *hopN1* | This study |
| *Psa3* V13 $\Delta CEL$ + p.*hopM1* (a.k.a. p.*shcM*<sub>*Psa*</sub>:*hopM1*) | *Psa3* V13 *CEL* knockout with plasmid-borne *shcM* (from *Psa3* V13) and *hopM1* (from *Psa3* V13) | This study |
| *Psa3* V13 $\Delta CEL$ + p.*avrE1* | *Psa3* V13 *CEL* knockout with plasmid-borne *avrE1* (lacking *shcE*) | This study |
| *Psa3* V13 $\Delta CEL$ + *hrpW1* | *Psa3* V13 *CEL* knockout with genomic knock-in of *hrpW1* | This study |
| *Psa3* V13 $\Delta CEL$ + *shcM:hopM1* | *Psa3* V13 *CEL* knockout with genomic knock-in of *shcM* and *hopM1* | This study |
| *Psa3* V13 $\Delta CEL$ + *shcE:avrE1* | *Psa3* V13 *CEL* knockout with genomic knock-in of *shcE* and *avrE1* | This study |
| *Pfo* Pf0-1 + p.EV | *P. fluorescens* (*Pfo*) Pf0-1 with empty vector | Jayaraman et al., 2017 |
| *Pfo* Pf0-1 + p.*hopN1* | *Pfo* Pf0-1 with plasmid-borne *shcN* and *hopN1* | Jayaraman et al., 2017 |
| *Pfo* Pf0-1 + p.*hopM1* (a.k.a. p.*shcM*<sub>*Psa*</sub>:*hopM1*) | *Pfo* Pf0-1 with plasmid-borne *shcM* (from *Psa3* V13) and *hopM1* | Jayaraman et al., 2017 |
| *Pfo* Pf0-1 + p.*avrE1* | *Pfo* Pf0-1 with plasmid-borne *avrE1* (lacking *shcE*) | Jayaraman et al., 2017 |
| *Pfo* Pf0-1 (T3S) + p.EV | *Pfo* Pf0-1 (T3S) with empty vector | Jayaraman et al., 2017 |
| *Pfo* Pf0-1 (T3S) + p.*hopN1* | *Pfo* Pf0-1 (T3S) with plasmid-borne *shcN* and *hopN1* | Jayaraman et al., 2017 |
| *Pfo* Pf0-1 (T3S) + p.*hopM1* (a.k.a. p.*shcM*<sub>*Psa*</sub>:*hopM1*) | *Pfo* Pf0-1 (T3S) with plasmid-borne *shcM* (from *Psa3* V13) and *hopM1* | Jayaraman et al., 2017 |
| *Pfo* Pf0-1 (T3S) + p.*avrE1* | *Pfo* Pf0-1 (T3S) with plasmid-borne *avrE1* (lacking *shcE*) | Jayaraman et al., 2017 |
| *Pfo* Pf0-1 + p.*shcM*<sub>*Pto*</sub>:*hopM1* | *Pfo* Pf0-1 with plasmid-borne *shcM* (from *Pto* DC3000) and *hopM1* (from *Psa3* V13) | This study |
| *Pfo* Pf0-1 (T3S) + p.*shcM*<sub>*Pto*</sub>:*hopM1* | *Pfo* Pf0-1 (T3S) with plasmid-borne *shcM* (from *Pto* DC3000) and *hopM1* (from *Psa3* V13) | This study |
| *Psa3* V13 $\Delta CEL$ | *Psa3* V13 *CEL* knockout with plasmid-borne *shcM* (from *Pto* DC3000) | This study |
| *Psa3* V13 $\Delta CEL$ + *shcM:hopM1* + p.*shcM*<sub>*Pto*</sub> | *Psa3* V13 *CEL* knockout with genomic knock-in of *shcM* and *hopM1* and plasmid-borne *shcM* (from *Pto* DC3000) | This study |
| *Psa3* V13 $\Delta CEL$ + *shcE:avrE1* + p.*shcM*<sub>*Pto*</sub> | *Psa3* V13 *CEL* knockout with genomic knock-in of *shcE* and *avrE1* and plasmid-borne *shcM* (from *Pto* DC3000) | This study |
| *Psa3* V13 $\Delta hopR1$ | *Psa3* V13 *hopR1* knockout | This study |
| *Psa3* V13 $\Delta hopR1$ + p.*hopR1* | *Psa3* V13 *hopR1* knockout with plasmid-borne *hopR1* | This study |
| *Psa3* V13 $\Delta CEL/\Delta hopR1$ | *Psa3* V13 *CEL* and *hopR1* double knockout | This study |
| *Psa3* V13 $\Delta hrcC$ + p.*shcM*<sub>*Psa*</sub>:*hopM1* | *Psa3* V13 *hrcC* knockout with plasmid-borne *shcM* (from *Psa3* V13) and *hopM1* (from *Psa3* V13) | This study |
| *Psa3* V13 $\Delta hrcC$ + p.*shcM*<sub>*Pto*</sub>:*hopM1* | *Psa3* V13 *hrcC* knockout with plasmid-borne *shcM* (from *Pto* DC3000) and *hopM1* (from *Psa3* V13) | This study |
| *Psa3* V13 $\Delta CEL$ + p.*shcM*<sub>*Pto*</sub>:*hopM1* | *Psa3* V13 *CEL* knockout with plasmid-borne *shcM* (from *Pto* DC3000) and *hopM1* (from *Psa3* V13) | This study |

**Supplementary Table S2.**
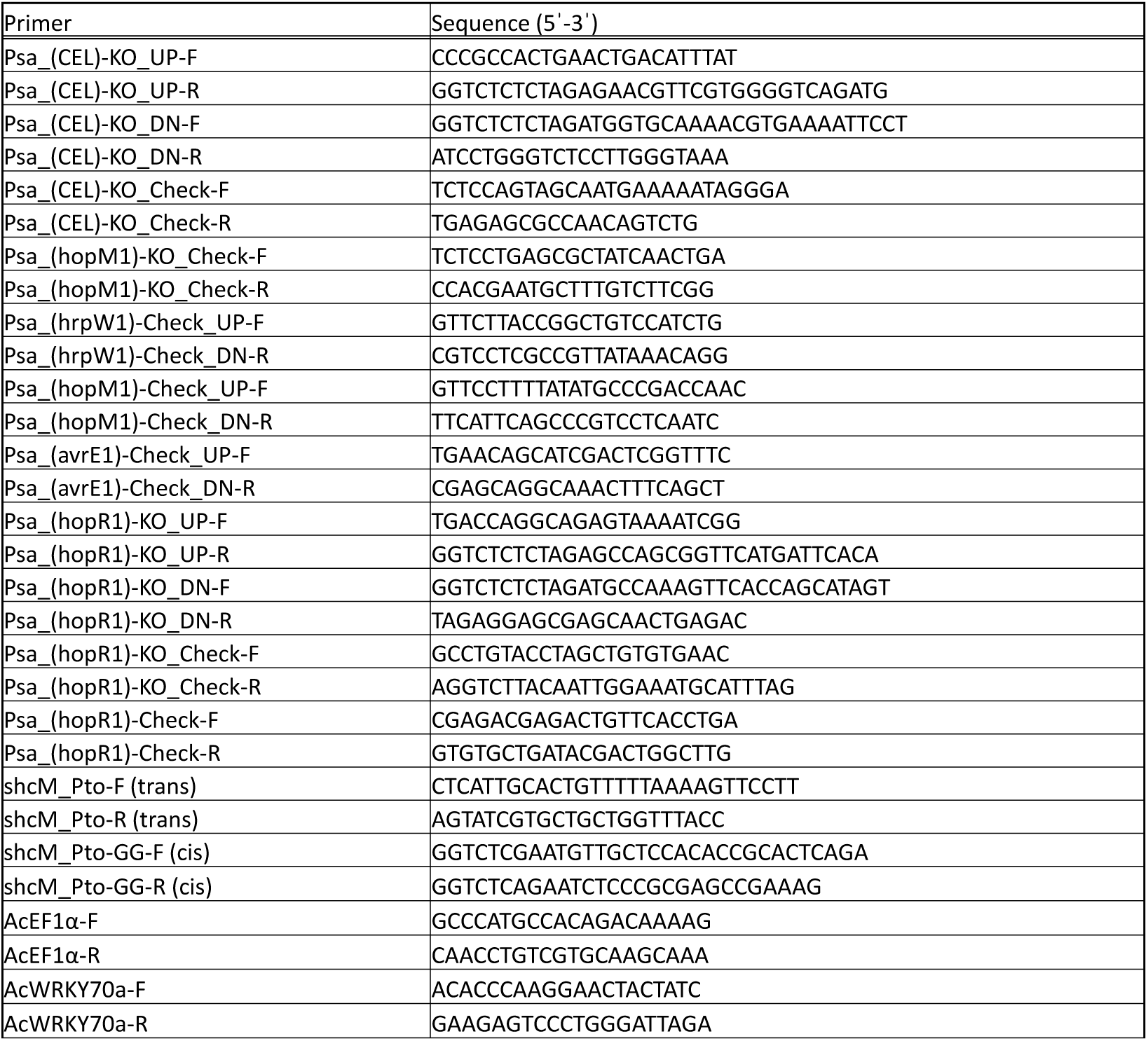

## References

Alfano, J.R., Charkowski, A.O., Deng, W.L., Badel, J.L., Petnicki-Ocwieja, T., van Dijk, K., Collmer, A., 2000. The *Pseudomonas syringae* Hrp pathogenicity island has a tripartite mosaic structure composed of a cluster of type III secretion genes bounded by exchangeable effector and conserved effector loci that contribute to parasitic fitness and pathogenicity in plants. Proc. Natl. Acad. Sci. U.S.A. 97, 4856–4861.

Badel, J.L., Nomura, K., Bandyopadhyay, S., Shimizu, R., Collmer, A., He, S.Y., 2003. *Pseudomonas syringae* pv. *tomato* DC3000 HopPtoM (CEL ORF3) is important for lesion formation but not growth in tomato and is secreted and translocated by the Hrp type III secretion system in a chaperone-dependent manner. Mol. Microbiol. 49, 1239–1251.

Badel, J.L., Shimizu, R., Oh, H.-S., Collmer, A., 2006. A *Pseudomonas syringae* pv. *tomato avrE1/hopM1* mutant is severely reduced in growth and lesion formation in tomato. Mol. Plant Microbe Interact. 19, 99–111. https://doi.org/10.1094/MPMI-19-0099

Baltrus, D.A., Nishimura, M.T., Romanchuk, A., Chang, J.H., Mukhtar, M.S., Cherkis, K., Roach, J., Grant, S.R., Jones, C.D., Dangl, J.L., 2011. Dynamic evolution of pathogenicity revealed by sequencing and comparative genomics of 19 *Pseudomonas syringae* isolates. PLoS Pathogens 7, e1002132. https://doi.org/10.1371/journal.ppat.1002132

Berge, O., Monteil, C.L., Bartoli, C., Chandeysson, C., Guilbaud, C., Sands, D.C., Morris, C.E., 2014. A user’s guide to a data base of the diversity of *Pseudomonas syringae* and its application to classifying strains in this phylogenetic complex. PLoS ONE 9, e105547. https://doi.org/10.1371/journal.pone.0105547

Büttner, D., He, S.Y., 2009. Type III protein secretion in plant pathogenic bacteria. Plant Physiol. 150, 1656–1664. https://doi.org/10.1104/pp.109.139089

Cai, R., Lewis, J., Yan, S., Liu, H., Clarke, C.R., Campanile, F., Almeida, N.F., Studholme, D.J., Lindeberg, M., Schneider, D., Zaccardelli, M., Setubal, J.C., Morales-Lizcano, N.P., Bernal, A., Coaker, G., Baker, C., Bender, C.L., Leman, S., Vinatzer, B.A., 2011. The plant pathogen *Pseudomonas syringae* pv. *tomato* is genetically monomorphic and under strong selection to evade tomato Immunity. PLoS Pathogens 7, e1002130. https://doi.org/10.1371/journal.ppat.1002130

Choi, K.-H., Kumar, A., Schweizer, H.P., 2006. A 10-min method for preparation of highly electrocompetent *Pseudomonas aeruginosa* cells: Application for DNA fragment transfer between chromosomes and plasmid transformation. Journal of Microbiological Methods 64, 391–397. https://doi.org/10.1016/j.mimet.2005.06.001

Choi, S., Jayaraman, J., Segonzac, C., Park, H.-J., Park, H., Han, S.-W., Sohn, K.H., 2017. *Pseudomonas syringae* pv. *actinidiae* type III effectors localized at multiple cellular compartments activate or suppress innate immune responses in *Nicotiana benthamiana*. Front Plant Sci 8, 2157. https://doi.org/10.3389/fpls.2017.02157

Clarke, C.R., Cai, R., Studholme, D.J., Guttman, D.S., Vinatzer, B.A., 2010. *Pseudomonas syringae* strains naturally lacking the classical *P. syringae hrp/hrc* locus are common leaf colonizers equipped with an atypical type III secretion system. MPMI 23, 198–210. https://doi.org/10.1094/MPMI-23-2-0198

Colombi, E., Straub, C., Künzel, S., Templeton, M.D., McCann, H.C., Rainey, P.B., 2017. Evolution of copper resistance in the kiwifruit pathogen *Pseudomonas syringae* pv. *actinidiae* through acquisition of integrative conjugative elements and plasmids. Environ. Microbiol. 19, 819–832. https://doi.org/10.1111/1462-2920.13662

Cunnac, S., Chakravarthy, S., Kvitko, B.H., Russell, A.B., Martin, G.B., Collmer, A., 2011. Genetic disassembly and combinatorial reassembly identify a minimal functional repertoire of type III effectors in *Pseudomonas syringae*. Proc. Natl. Acad. Sci. U.S.A. 108, 2975–2980. https://doi.org/10.1073/pnas.1013031108

Deb, D., Mackey, D., Opiyo, S.O., McDowell, J.M., 2018. Application of alignment-free bioinformatics methods to identify an oomycete protein with structural and functional similarity to the bacterial AvrE effector protein. PLoS ONE 13, e0195559. https://doi.org/10.1371/journal.pone.0195559

DebRoy, S., Thilmony, R., Kwack, Y.-B., Nomura, K., He, S.Y., 2004. A family of conserved bacterial effectors inhibits salicylic acid-mediated basal immunity and promotes disease necrosis in plants. Proceedings of the National Academy of Sciences 101, 9927–9932. https://doi.org/10.1073/pnas.0401601101

Degrave, A., Siamer, S., Boureau, T., Barny, M.-A., 2015. The AvrE superfamily: ancestral type III effectors involved in suppression of pathogen-associated molecular pattern-triggered immunity. Molecular Plant Pathology 16, 899–905. https://doi.org/10.1111/mpp.12237

Deng, W.-L., Rehm, A.H., Charkowski, A.O., Rojas, C.M., Collmer, A., 2003. *Pseudomonas syringae* exchangeable effector loci: Sequence diversity in representative pathovars and virulence function in *P. syringae* pv. *syringae* B728a. Journal of Bacteriology 185, 2592–2602. https://doi.org/10.1128/JB.185.8.2592-2602.2003

Dillon, M.M., Almeida, R.N.D., Laflamme, B., Martel, A., Weir, B.S., Desveaux, D., Guttman, D.S., 2019a. Molecular evolution of *Pseudomonas syringae* type III secreted effector proteins. Front. Plant Sci. 10, 418. https://doi.org/10.3389/fpls.2019.00418

Dillon, M.M., Thakur, S., Almeida, R.N.D., Wang, P.W., Weir, B.S., Guttman, D.S., 2019b. Recombination of ecologically and evolutionarily significant loci maintains genetic cohesion in the *Pseudomonas syringae* species complex. Genome Biology 20. https://doi.org/10.1186/s13059-018-1606-y

Engler, C., Kandzia, R., Marillonnet, S., 2008. A one pot, one step, precision cloning method with high throughput capability. PLoS ONE 3, e3647. https://doi.org/10.1371/journal.pone.0003647

Huynh, T., Dahlbeck, D., Staskawicz, B., 1989. Bacterial blight of soybean: regulation of a pathogen gene determining host cultivar specificity. Science 245, 1374–1377. https://doi.org/10.1126/science.2781284

Jayaraman, J., Choi, S., Prokchorchik, M., Choi, D.S., Spiandore, A., Rikkerink, E.H., Templeton, M.D., Segonzac, C., Sohn, K.H., 2017. A bacterial acetyltransferase triggers immunity in *Arabidopsis thaliana* independent of hypersensitive response. Sci Rep 7, 3557. https://doi.org/10.1038/s41598-017-03704-x

Jin, L., Ham, J.H., Hage, R., Zhao, W., Soto-Hernández, J., Lee, S.Y., Paek, S.-M., Kim, M.G., Boone, C., Coplin, D.L., Mackey, D., 2016. Direct and indirect targeting of PP2A by conserved bacterial type-III effector proteins. PLoS Pathog. 12, e1005609. https://doi.org/10.1371/journal.ppat.1005609

Karasov, T.L., Almario, J., Friedemann, C., Ding, W., Giolai, M., Heavens, D., Kersten, S., Lundberg, D.S., Neumann, M., Regalado, J., Neher, R.A., Kemen, E., Weigel, D., 2018. *Arabidopsis thaliana* and *Pseudomonas* pathogens exhibit stable associations over evolutionary timescales. Cell Host & Microbe 24, 168-179.e4. https://doi.org/10.1016/j.chom.2018.06.011

Kovach, M.E., Elzer, P.H., Steven Hill, D., Robertson, G.T., Farris, M.A., Roop, R.M., Peterson, K.M., 1995. Four new derivatives of the broad-host-range cloning vector pBBR1MCS, carrying different antibiotic-resistance cassettes. Gene 166, 175–176. https://doi.org/10.1016/0378-1119(95)00584-1

Kvitko, B.H., Park, D.H., Velásquez, A.C., Wei, C.-F., Russell, A.B., Martin, G.B., Schneider, D.J., Collmer, A., 2009. Deletions in the repertoire of *Pseudomonas syringae* pv. *tomato* DC3000 type III secretion effector genes reveal functional overlap among effectors. PLoS Pathogens 5, e1000388. https://doi.org/10.1371/journal.ppat.1000388

Kvitko, B.H., Ramos, A.R., Morello, J.E., Oh, H.-S., Collmer, A., 2007. Identification of harpins in *Pseudomonas syringae* pv. *tomato* DC3000, which are functionally similar to HrpK1 in promoting translocation of type III secretion system effectors. Journal of Bacteriology 189, 8059–8072. https://doi.org/10.1128/JB.01146-07

Laflamme, B., Dillon, M.M., Martel, A., Almeida, R.N.D., Desveaux, D., Guttman, D.S., 2020. The pan-genome effector-triggered immunity landscape of a host-pathogen interaction. Science 367, 763–768. https://doi.org/10.1126/science.aax4079

Lorang, J.M., Keen, N.T., 1995. Characterization of *avrE* from *Pseudomonas syringae* pv. *tomato*: a *hrp*-linked avirulence locus consisting of at least two transcriptional units. Mol. Plant Microbe Interact. 8, 49–57.

McAtee, P.A., Brian, L., Curran, B., van der Linden, O., Nieuwenhuizen, N.J., Chen, X., Henry-Kirk, R.A., Stroud, E.A., Nardozza, S., Jayaraman, J., Rikkerink, E.H.A., Print, C.G., Allan, A.C., Templeton, M.D., 2018. Re-programming of *Pseudomonas syringae* pv. *actinidiae* gene expression during early stages of infection of kiwifruit. BMC Genomics 19. https://doi.org/10.1186/s12864-018-5197-5

McCann, H.C., Li, L., Liu, Y., Li, D., Pan, H., Zhong, C., Rikkerink, E.H.A., Templeton, M.D., Straub, C., Colombi, E., Rainey, P.B., Huang, H., 2017. Origin and evolution of the kiwifruit canker pandemic. Genome Biology and Evolution 9, 932–944. https://doi.org/10.1093/gbe/evx055

Mesarich, C.H., Rees-George, J., Gardner, P.P., Ghomi, F.A., Gerth, M.L., Andersen, M.T., Rikkerink, E.H.A., Fineran, P.C., Templeton, M.D., 2017. Transposon insertion libraries for the characterization of mutants from the kiwifruit pathogen *Pseudomonas syringae* pv. *actinidiae*. PLoS ONE 12, e0172790. https://doi.org/10.1371/journal.pone.0172790

Monteil, C.L., Cai, R., Liu, H., Mechan Llontop, M.E., Leman, S., Studholme, D.J., Morris, C.E., Vinatzer, B.A., 2013. Nonagricultural reservoirs contribute to emergence and evolution of *Pseudomonas syringae* crop pathogens. New Phytologist 199, 800–811. https://doi.org/10.1111/nph.12316

Morris, C.E., Lamichhane, J.R., Nikolic, I., Stankovic, S., Moury, B., 2019. The overlapping continuum of host range among strains in the *Pseudomonas syringae* complex. Phytopathol Res 1, 4. https://doi.org/10.1186/s42483-018-0010-6

Munkvold, K.R., Russell, A.B., Kvitko, B.H., Collmer, A., 2009. *Pseudomonas syringae* pv. *tomato* DC3000 type III effector HopAA1-1 functions redundantly with chlorosis-promoting factor PSPTO4723 to produce bacterial speck lesions in host tomato. MPMI 22, 1341–1355. https://doi.org/10.1094/MPMI-22-11-1341

Nomura, K., Mecey, C., Lee, Y.-N., Imboden, L.A., Chang, J.H., He, S.Y., 2011. Effector-triggered immunity blocks pathogen degradation of an immunity-associated vesicle traffic regulator in Arabidopsis. Proceedings of the National Academy of Sciences 108, 10774–10779. https://doi.org/10.1073/pnas.1103338108

Poulter, R.T.M., Ho, J., Handley, T., Taiaroa, G., Butler, M.I., 2018. Comparison between complete genomes of an isolate of *Pseudomonas syringae* pv. actinidiae from Japan and a New Zealand isolate of the pandemic lineage. Sci Rep 8, 10915. https://doi.org/10.1038/s41598-018-29261-5

Roine, E., Wei, W., Yuan, J., Nurmiaho-Lassila, E.-L., Kalkkinen, N., Romantschuk, M., He, S.Y., 1997. Hrp pilus: An *hrp*-dependent bacterial surface appendage produced by *Pseudomonas syringae* pv. *tomato* DC3000. Proceedings of the National Academy of Sciences 94, 3459–3464. https://doi.org/10.1073/pnas.94.7.3459

Schäfer, A., Tauch, A., Jäger, W., Kalinowski, J., Thierbach, G., Pühler, A., 1994. Small mobilizable multi-purpose cloning vectors derived from the Escherichia coli plasmids pK18 and pK19: selection of defined deletions in the chromosome of *Corynebacterium glutamicum*. Gene 145, 69–73.

Templeton, M.D., Warren, B.A., Andersen, M.T., Rikkerink, E.H.A., Fineran, P.C., 2015. Complete DNA sequence of *Pseudomonas syringae* pv. *actinidiae*, the causal agent of kiwifruit canker disease. Genome Announc 3. https://doi.org/10.1128/genomeA.01054-15

Thomas, W.J., Thireault, C.A., Kimbrel, J.A., Chang, J.H., 2009. Recombineering and stable integration of the *Pseudomonas syringae* pv. *syringae* 61 *hrp/hrc* cluster into the genome of the soil bacterium *Pseudomonas fluorescens* Pf0-1: Stable integration of a T3SS-locus into Pf0-1. The Plant Journal 60, 919–928. https://doi.org/10.1111/j.1365-313X.2009.03998.x

Weber, E., Engler, C., Gruetzner, R., Werner, S., Marillonnet, S., 2011. A modular cloning system for standardized assembly of multigene constructs. PLoS ONE 6, e16765. https://doi.org/10.1371/journal.pone.0016765

Wei, C.-F., Kvitko, B.H., Shimizu, R., Crabill, E., Alfano, J.R., Lin, N.-C., Martin, G.B., Huang, H.-C., Collmer, A., 2007. A *Pseudomonas syringae* pv. *tomato* DC3000 mutant lacking the type III effector HopQ1-1 is able to cause disease in the model plant *Nicotiana benthamiana*. The Plant Journal 51, 32–46. https://doi.org/10.1111/j.1365-313X.2007.03126.x

Wei, H.-L., Zhang, W., Collmer, A., 2018. Modular study of the type III effector repertoire in *Pseudomonas syringae* pv. *tomato* DC3000 reveals a matrix of effector interplay in pathogenesis. Cell Reports 23, 1630–1638. https://doi.org/10.1016/j.celrep.2018.04.037

Worley, J.N., Russell, A.B., Wexler, A.G., Bronstein, P.A., Kvitko, B.H., Krasnoff, S.B., Munkvold, K.R., Swingle, B., Gibson, D.M., Collmer, A., 2013. *Pseudomonas syringae* pv. *tomato* DC3000 CmaL (PSPTO4723), a DUF1330 family member, is needed to produce L-*allo*-Isoleucine, a precursor for the phytotoxin coronatine. J. Bacteriol. 195, 287–296. https://doi.org/10.1128/JB.01352-12

Xin, X.-F., Kvitko, B., He, S.Y., 2018. *Pseudomonas syringae*: what it takes to be a pathogen. Nature Reviews Microbiology 16, 316–328. https://doi.org/10.1038/nrmicro.2018.17

Xin, X.-F., Nomura, K., Aung, K., Velásquez, A.C., Yao, J., Boutrot, F., Chang, J.H., Zipfel, C., He, S.Y., 2016. Bacteria establish an aqueous living space in plants crucial for virulence. Nature 539, 524–529. https://doi.org/10.1038/nature20166

Zembek, P., Danilecka, A., Hoser, R., Eschen-Lippold, L., Benicka, M., Grech-Baran, M., Rymaszewski, W., Barymow-Filoniuk, I., Morgiewicz, K., Kwiatkowski, J., Piechocki, M., Poznanski, J., Lee, J., Hennig, J., Krzymowska, M., 2018. Two strategies of *Pseudomonas syringae* to avoid recognition of the HopQ1 effector in *Nicotiana* species. Front. Plant Sci. 9, 978. https://doi.org/10.3389/fpls.2018.00978

